# The role of *Akkermansia muciniphila* sulfatases in colonic mucin utilisation

**DOI:** 10.1101/2025.09.11.675649

**Authors:** Debajit Dey, Naba D. Salman, Charles W.E. Tomlinson, Chunsheng Jin, Grete Raba, Sadie R. Schaus, Marcus Nilsson, Zak McIver, Adam Simpkin, Matthew Davy, Daniel J Rigden, Mirjam Czjzek, Dominic P Byrne, Mihai Oltean, Alexander Case, Christoph G. Baumann, Gareth SA Wright, Sjoerd van der Post, Edwin A. Yates, Eric C. Martens, Lauren Davey, Ana S. Luis, Alan Cartmell

**Affiliations:** Department of Biology, University of York, Heslington, York, YO10 5DD, UK; Department of Chemistry, York Structural Biology Laboratory (YSBL), University of York, Wentworth Way, York, YO10 5DD, UK; York Biomedical Research Institute (YBRI), University of York, Wentworth Way, York, YO10 5DD, UK; Department of Medical Biochemistry and Cell Biology, Institute of Biomedicine, University of Gothenburg, 40530, Gothenburg, Sweden; Proteomics Core Facility at Sahlgrenska Academy, University of Gothenburg, 40530, Gothenburg, Sweden; Department of Chemistry and Biotechnology, Tallinn University of Technology, 12618, Tallinn, Estonia; Department of Microbiology and Immunology, University of Michigan, Ann Arbor, MI 48109, USA; Department of Biochemistry, Cell and Systems Biology, Institute of Systems, Molecular and Integrative Biology, University of Liverpool, Liverpool L69 7ZB, UK; Sorbonne Université, Univ Paris 06, CNRS, UMR 8227, Integrative Biology of Marine Models, Station Biologique de Roscoff, CS 90074, Roscoff, Bretagne, France; Department of Transplantation Surgery, Institute for Clinical Sciences, Sahlgrenska Academy, University of Gothenburg, 41345, Gothenburg, Sweden; Transplant Institute, Sahlgrenska University Hospital, 41345, Gothenburg, Sweden; School of Life Sciences, University of Essex, Wivenhoe Park, Colchester, CO4 3SQ, UK; Department of Biochemistry and Microbiology, University of Victoria, Victoria, British Columbia, Canada; SciLifeLab, University of Gothenburg, 41390 Gothenburg, Sweden

## Abstract

*Akkermansia muciniphila*, an obligate mucin degrader, is a major member of the human colonic microbiota and has been associated positive health outcomes. Mucins are complex glycoproteins that contain heavily sulfated *O*-glycans and form the protective colonic mucus layer. Bacterial carbohydrate sulfatases are required to metabolise these heavily sulfated mucin glycans and excessive bacterial foraging has been associated with several diseases. Sulfatases have been linked with inflammatory bowel disease, making these microbiota enzymes potential drug targets. *A. muciniphila* expresses carbohydrate sulfatases that can act on colonic mucins yet their roles in its metabolism remain opaque. Our data reveal that *A. muciniphila* requires glycopeptides/protein forms of colonic mucin for metabolism and its sulfatases have unique adaptations compared to *Bacteroides* species. Localisation studies reveal that desulfation of *N*-acetyl-D-glucosamine, but not D-galactose, is exclusively periplasmic. A cell surface sulfatase has a novel carbohydrate binding module that binds to colonic mucin. This paints a contrasting picture of sulfated mucin metabolism by *Akkermansia muciniphila* versus *Bacteroides* species. These data will be important for understanding the contexts for *Akkermansia muciniphila*’s positive health correlations.

## Introduction

The human colonic microbiota (HCM) is a vast microbial community composed of hundreds of bacterial species that greatly impact the host’s physiology^1,2^. Colonocytes, the cells lining the colon, derive ∼70 % of their energy content from short chain fatty acids^3^ (SCFAs) produced by HCM carbohydrate fermentation^4^. These SCFAs have been shown to have anti-inflammatory^5^ and anti-cancer effects^6^, and implications in degenerative brain disease^7,8^.

In the colon, the host’s mucins form a protective barrier that separates the HCM from the host epithelial layer but also serve as both a nutrient source and colonisation factor for the HCM^4,9^. MUC2 is the major gel-forming colonic mucin and up to 80 % of its mass can be *O*-glycans^10^. Sulfation of mucin *O*-glycans is variable along the gastrointestinal tract increasing from proximal to distal regions with colonic MUC2 having the highest sulfation level^11^ (up to 10 % by mass)^12^. This sulfation adds complexity and requires colonic bacteria to express carbohydrate sulfatases to fully metabolise colonic mucin^9,13^. All known carbohydrate sulfatases belong to the S1 family of sulfatases which is divided into 110 subfamilies denoted as S1_X^14,15^.

A physiologically healthy mucosal barrier must be maintained for HCM eubiosis^16,17^. Increased activity of HCM carbohydrate-active enzymes can lead to an excessive degradation of the mucin *O*-glycans^18,19^, disruption of the protective mucus barrier, and diseases such as ulcerative colitis (UC)^20,21^; a type of inflammatory bowel disease. Indeed, some HCM species such as *Bacteroides thetaiotaomicron* (*B. theta*) and *Bacteroides vulgatus* have been shown to drive UC in a sulfatase-dependent manner in a susceptible mice model^21,22^. HCM carbohydrate sulfatase activity is also increased in humans with active UC^19^. By contrast, the mucin-specialist *Akkermansia muciniphila* (*A. muc*) that can account for up to ∼3 % of the total HCM^23,24^ encodes multiple carbohydrate sulfatases, but its prevalence in human disease association studies was inversely proportional to disease and inflammation markers^25–27^. Whilst, in animal models its beneficial effects were context dependent. On low fibre diets the presence of *A. muc* increased pathogen susceptibility^28^ and food driven allergy^29^ but, on high fibre diet these effects were nullified with *A. muc* linked to a reduced pathogen susceptibility^28^. Finally, Increased *A. muc* abundance in Il10 deficient mice, via repeated oral gavage, was sufficient to drive colitis-like disease^30^.

The *A. muc* genome encodes 10 S1 carbohydrate sulfatases, classified into 6 subfamilies, that are related to the mucin targeting sulfatases described in *B. theta*^9,31^. To date most studies of *A. muc*, focused on characterising its glycoside hydrolases (GHs), have used gastric mucins; substrates low in sulfation and only on the *O*6 position of *N*-acetyl-D-glucosamine. Recent data, however, has shown that for heavily sulfated colonic mucin glycoprotein, and not free colonic mucin oligosaccharides (cMOs), are needed to support growth^32,33^. Yet, no detailed work on the *A. muc* carbohydrate sulfatases, and their role in metabolising heavily sulfated colonic mucins, has been conducted. Recently, limited analysis of *A. muc* sulfatase activity was performed on simple oligosaccharides, but this study did not reveal detailed activity on colonic *O*-glycans, kinetic, structural, or cellular location information^34^. Such knowledge is critical to understand the role of *A. muc* carbohydrate sulfatases in the metabolism of heavily sulfated colonic mucin.

In this work we used wildtype and sulfatase transposon mutants to analyse *A. muc* growth on colonic mucins revealing processed glycoproteins fragments, not colonic mucin glycoproteins, are *A. muc’s* preferred substrate. We provide detailed kinetic information for 8 of the 10 *A. muc* S1 sulfatases, define their cellular location, and structurally characterize 5 of these enzymes. We also identify the first sulfatase linked carbohydrate-binding module which binds colonic mucin and is restricted to *Akkermansia* species. These data reveal significant differences in substrate specificity, sulfatase enzyme structure and modularity, and enzyme location compared to mucin degrading *Bacteroides* species. This opens the potential to develop disease targeting *Bacteroides* sulfatase drugs that spare *A. muc* in contexts were its abundance is health positive, whilst also informing the sulfobiology of this important HCM member.

## Results

### A. muciniphila requires glycoprotein forms of colonic glycans for growth

We initially established growth profiles of *A. muc* on soluble porcine gastric mucin (sPGM) and gastric mucin oligosaccharides (gMOs) released from sPGM glycoprotein. The data showed robust growth on sPGM **(Supplementary figure 1a)**. Growth on gMOs by *A. muc* was much weaker showing a gradual increase in growth and reaching a lower level than on sPGM, by contrast *B. theta*, and a mutant unable to use contaminating glycosaminoglycans (*B. theta*^ΔCS/HS^), grew robustly on gMOs **(Figure 1a)**. We next examined growth on a soluble protein fraction extracted from PCM containing only protein (∼5.0 % MUC2 fragments and ∼95 % cellular protein from cell lysis; **Supplementary dataset 1**) and colonic mucin oligosaccharides (cMOs). *A. muc* was incapable of growth on these substrates whilst *B. theta* growth was supported by cMOs^9^ **(Figure 1a and Supplementary figure 1a)**. After the preparation of a new batch of cMOs we observed strong growth for *A. muciniphila*, whilst *B. theta* was also able to utilise this substrate but to a lower degree **(Figure 1a)**. We hypothesized that differences in available glycans between the two substrate batches would be responsible for the differences in growth profile. Comparison of the substrates, by NMR **(Figure 1b and supplementary figure 1b)**, glycoprotein gel staining **(Figure 1c)**, and mass spectrometry **(Figure 1d and Supplementary dataset 2)** revealed that the second batch of cMOs is enriched in glycopeptides/proteins compared to initial cMO batch and their NMR profile was more similar to sPGM. For this reason cMO batch 2 is termed glycopeptide/glycoprotein colonic mucin oligosaccharide (gpcMOs). Following this we treated porcine colonic mucin (PCM) with the proteases trypsin and proteinase K to generate soluble mucin glycoproteins. These hydrolysed substrates demonstrated similar NMR profiles to gpcMOs and sPGM **(Supplementary figure 1b)** but supported growth of *A. muc* to lowers levels **(Supplementary figure 1a),** with growth on trypsin treated material requiring a doubled substrate concentration for robust growth. These data reveal that *A. muc* targets glycopeptide/protein forms of mucin and these are needed for robust growth on gastric mucins but absolutely required for growth on colonic mucins. By contrast, *B. theta* could grow on gMOs, cMOs, and gpcMOs but grew more poorly on gpcMOs **(Figure 1a)**. This indicates that *A. muc* targets specific glycopeptides of a particular size and/or motif that were serendipitously captured in our gpcMO preparation, and these cannot be fully recapitulated by trypsin or proteinase K treatment.

**Figure 1.**
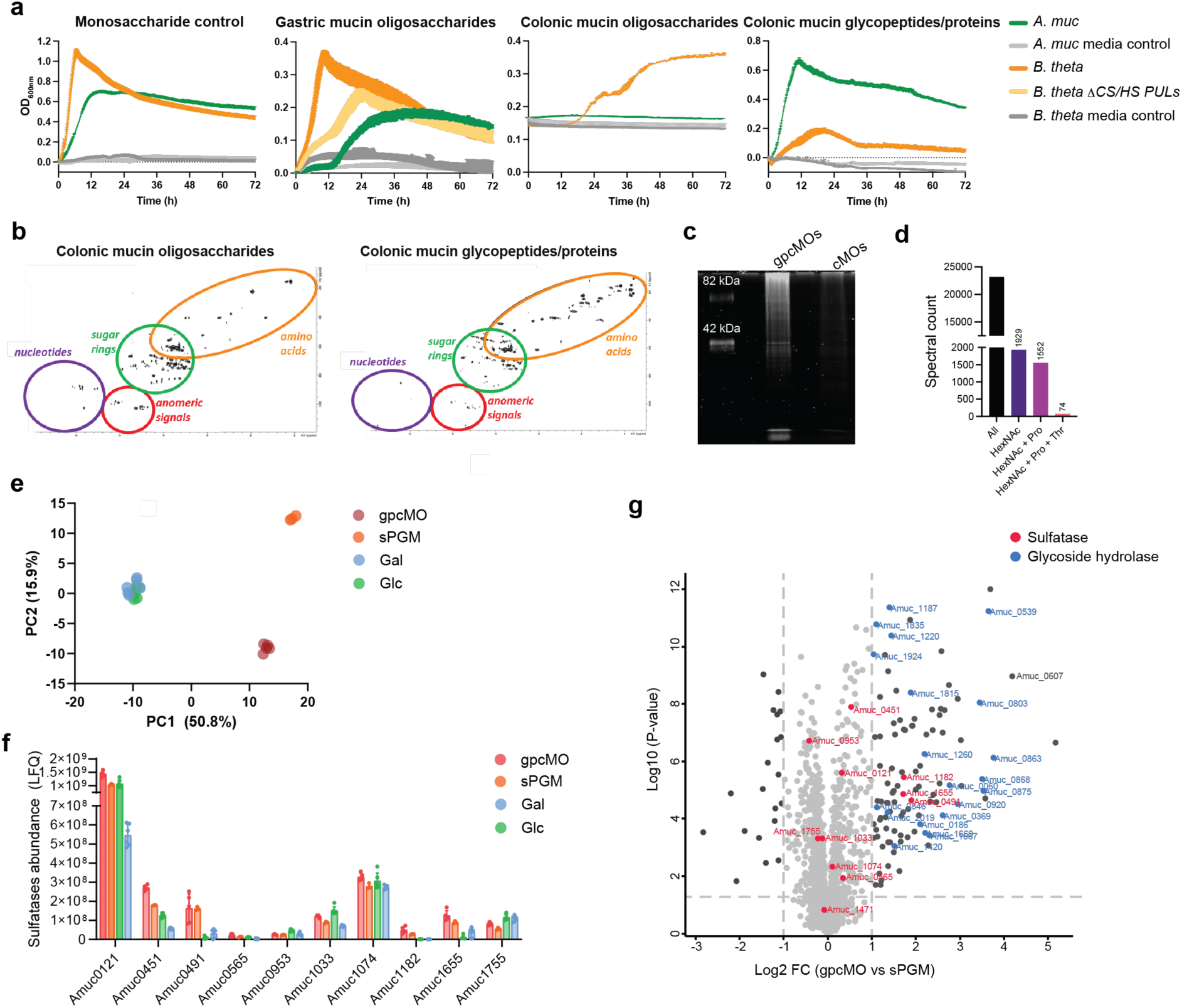
Growth profiles and protein expression of *A. muciniphila* on variable mucin structures. **a.** Growth profiles of *A. muciniphila* on low sulfation and high sulfation mucin *O*-glycans and glycopeptide/proteins. All substrate concentrations are 5 mg/ml except GlcNAc which was used at 10 mg/ml for *A. muciniphila* whilst glucose was used for *B. theta* in the monosaccharide controls. **b.** left, NMR profiles of porcine colonic mucin oligosaccharides and porcine colonic glycopeptides/proteins **c.** glycoprotein gel staining of porcine colonic mucin oligosaccharides and porcine colonic glycopeptides/proteins. **d.** Spectral counts of glycopeptides and glycan-containing spectra identified in the porcine colonic glycoprotein fraction by mass spectrometry. **e.** Principal component analysis of global proteomic data **f.** Total abundance of *A. muciniphila* sulfatases across proteomic samples on from *A. muciniphila* grown on various carbon sources **g.** Proteomic analysis of relative protein expression of *A. muciniphila* grown on glucose or gpcMOs as a sole carbon source.

To determine if the increased sulfation present in colonic mucin drove increased expression of *A. muc* carbohydrate sulfatases we performed proteomic analysis of *A. muc* grown on D-glucose (Glc), D-galactose (Gal), sPGM and gpcMOs. The data showed that the mucin samples cluster differently from each other and from the monosaccharides (**Figure 1e**). All 10 *A. muc* sulfatases were detected in all conditions with only three sulfatases (Amuc0491, Amuc1655 and Amuc1182) significantly upregulated by mucin glycans (**Figure 1f and g**). Overall, the different gastric and colonic mucin substrates did not lead to large differences in sulfatase expression **(Supplemental figure 1c and Supplementary dataset 3).** Although, in presence of gpcMO the expression of Amuc_0627, a glycoprotease associated with mucin degradation, was significantly upregulated by gpcMOs compared to sPGM **(Figure 1g)**. This suggests that the protease activity is required to the utilization of the glycoprotein present in the substrate.

*A. muciniphila sulfatases effectively desulfate model and colonic oligosaccharide substrates* To understand the biochemistry of how *A. muc* desulfates mucin *O*-glycans we expressed all ten S1 sulfatases recombinantly and detected activities for 8 of the 10 enzymes **(Figure 2a and Supplementary figure 3)**. Only Amuc_0565 and Amuc_1182, from subfamilies S1_4 and S1_19 displayed no activities against any substrates tested. This is in line with a recently published qualitative study^34^. To define the true substrate specificity of each sulfatase we determined the catalytic efficiency and pH optimum on fluorescently labelled substrates **(Supplementary figure 2 and Supplementary table 1)** as well as activity on model and mucin derived glycans **(Figure 2a,b,c and Supplementary figure 3)**.

**Figure 2.**
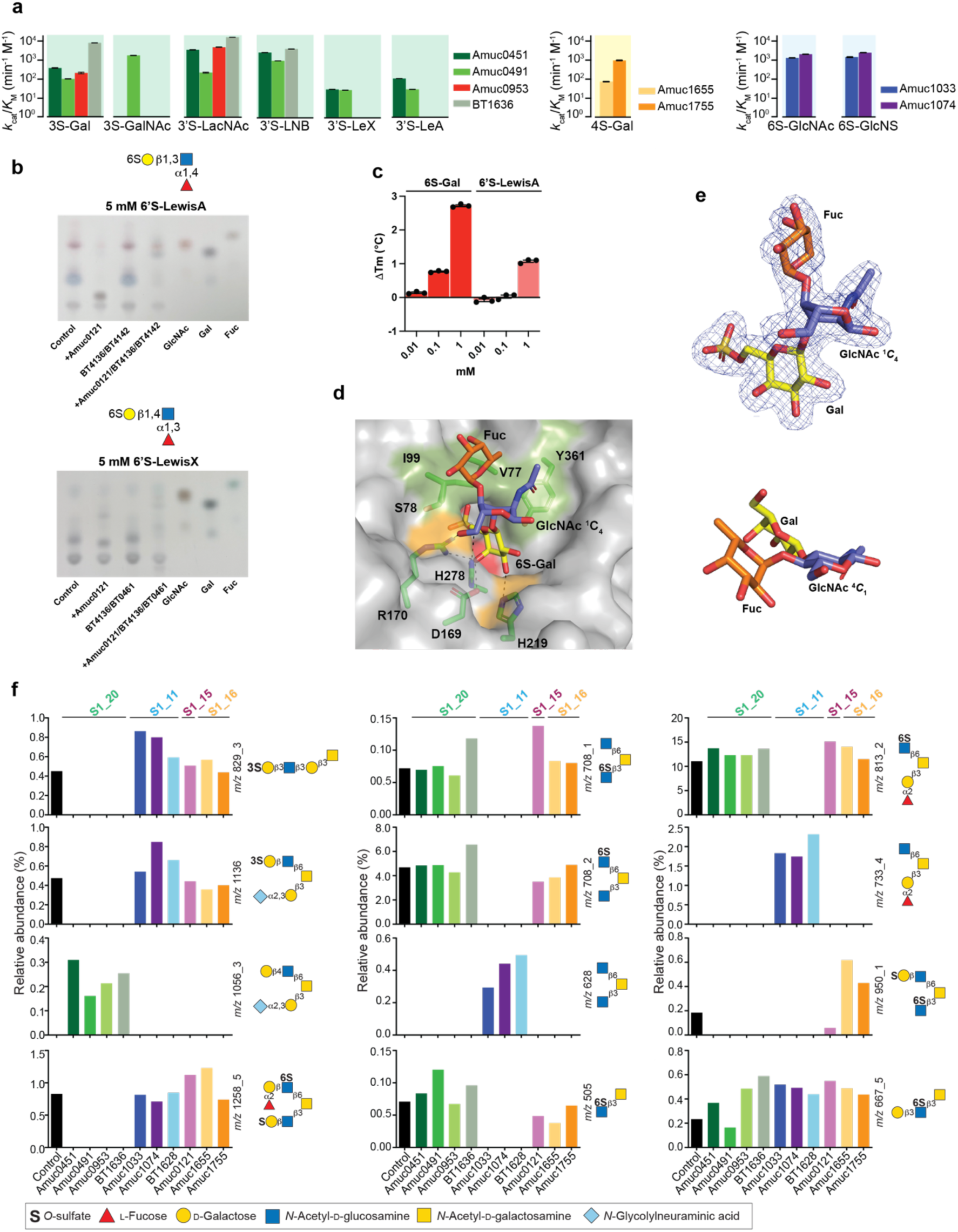
Substrate preference and activity of S1 *A. muciniphila* sulfatases. **a.** Kinetic activity for sulfatases for which an activity could be identified. Assays were carried out in an appropriate buffer for the pH optimum for each respective enzyme with 1 μM of a fluorescently labelled substrate, 150 mM NaCl, and 5 mM CaCl_2_. Experiments are technical triplicates. **b.** Thin layer chromatography demonstrating Amuc0121^6S-Gal^ is essential to breakdown *O*6 Gal sulfated LewisA and LewisX antigen. BT4136 is a GH95 fucosidase capable of acting on α1,3/4 linkage whilst BT0461 and BT4241 are a GH2 and GH109 galactosidases targeting β1,4, and β1,3 glycosidic linkages; these enzymes were at a final concentration of 1 μM. The final concentration of Amuc0121^6S-Gal^ was 100 μM. Assays were carried out in 10 mM HEPES, pH 7.0. with 150 mM NaCl at 37°C overnight. **c.** Thermostability shift assay suggesting 6S-Gal binds more favourably than 6’S-LewisA. **d.** Surface representation of the Amuc0121^6S-Gal^ crystal structure in complex with 6’S-LewisA with key residue highlighted beneath. In orange is the galactose recognition triad whilst green highlights the hydrophobic patch that accommodate Fuc (L-fucose); 6S-Gal is O6 sulfated D-galactose whilst GlcNAc is N-acetyl-D-glucosamine. **e.** The weighted 2mFobs-DFc electron density map, contoured at 1 σ, for the 6’S-LewisA substrate from **d.** The GlcNAc adopts a strained ^1^*C*_4_ conformation. Below this is a relaxed LewisA substrate with the GlcNAc in a ^4^*C*_1_ conformation from PDB: 6ORF.**f** Relative abundance of processed glycan species generated by activity of *A. muciniphila* sulfatases on colonic mucin oligosaccharides, as assessed by mass spectrometry.

*A. muc* contains 3 S1_20 sulfatases Amuc0451^3S-LacNAc^, Amuc0491^3S-Gal/GalNAc^, and Amuc0953^3S-LacNAc^ that desulfated *O*3 sulfated D-galactose (3S-Gal) containing substrates **(Figure 2a and Supplementary figure 3)** although only Amuc0491^3S-Gal/GalNAc^ was capable of desulfating *O*3 sulfated *N*-acetyl D-galactose (3S-GalNAc) demonstrating a ∼10 fold preference for this substrate **(Figure 2a and Supplementary figure 3)**. Amuc0451^3S-Gal^ and Amuc0953^3S-Gal^ desulfated 3S-Gal, *O*3 sulfated lactosamine (3’S-LacNAc), *O*3 sulfated lacto-*N*-biose (3’S-LNB), and both 3’S-Lewis X and 3’S-Lewis A but were ∼10 fold more active on 3’S-LacNAc and 3’S-LNB than 3S-Gal **(Figure 2a)**.

A single S1 sulfatase, Amuc0121^6S-Gal/GalNAc^, from S1_15 was found to desulfate both *O*6 sulfated D-galactose (6S-Gal) and *O*6 sulfated *N*-acetyl D-galactose (6S-GalNAc) **(Supplementary figure 3)**, as well as the trisaccharides 6’S-LewisA and 6’S-LewisX **(Figure 2b)**, and is the only enzyme known to breakdown these Lewis structures, although its affinity for the 6’S-LewisA is lower than for 6S-Gal **(Figure 2c).** Crystal structures of Amuc0121^6S-Gal/GalNAc^ in complex with Gal, 6S-Gal, and 6’S-LewisA reveal that Gal is bound identically in all three with *O*1 pointing into solvent **(Figure 2d and Supplementary figure 4a,b)**. Surprisingly, the binding of 6’S-LewisA requires the GlcNAc to adopt an atypical, strained, ^1^*C*_4_ chair conformation instead of the lower energy ^4^*C*_1_ **(Figure 2e)**. This allows the Fuc to sit above a hydrophobic patch created by the conserved residues Val77, Ile99, and Tyr361^31^, whilst the *O*6 of *N*-acetyl-D-glucosamine (GlcNAc) interacts with the conserved residue Arg170 that is part of the Gal recognition triad within roughly half of S1_15 members^31^. No kinetic measurement could be accurately made for the Amuc0121^6S-Gal/GalNAc^ likely due to lower FGly installation by *E. coli* during recombinant expression, reported previously for the S1_15 subfamily^31^.

The two S1_16 sulfatases, Amuc1655^4S-Gal^ and Amuc1755^4S-Gal^, were shown to have specificity for *O*4 sulfated D-galactose (4S-Gal) with Amuc1755^4S-Gal^ appearing to be ∼10 fold more efficient than Amuc1655^4S-Gal^ **(Figure 2a)**. Although both enzymes show very weak desulfation of *O*4 sulfated *N*-acetyl D-galactose (4S-GalNAc) **(Supplementary figure 3)**. The final two characterised enzymes were Amuc1033^6S-GlcNAc^ and Amuc1074^6S-GlcNAc^. The enzymes belong to S1_11 and were able to desulfate both 6S-GlcNAc and *O*6 sulfated *N*-sulfo D-glucsoamine BODIPY labelled substrates with equal efficiency **(Figure 2a).**

Having determined activities on model substrates we next incubated these enzymes with cMOs and observed the effect on the relative abundance of each glycan by mass spectrometry **(Figure 2f and Supplementary dataset 4**). Only the 3S-Gal/GalNAc and 6S-GlcNAc sulfatases were active on these cMOs. No difference in activity was observed between the three S1_20 or the two S1_11 enzymes and all the five sulfatases were only active on terminal sulfated linkages **(Figure 2f)**. These results are similar to the previously reported activities for *B. theta* sulfatases^9^ and indicate that *A. muc* encodes sulfatases to remove the main sulfated linkages found in complex cMOs.

### A. muciniphila restricts N-acetyl-D-glucosamine desulfation to the periplasm

To reveal the cellular localisation of *A. muc* sulfatases we performed assays using whole cells and cell free extracts (CFE) using wildtype bacteria and sulfatase transposon mutants grown on GlcNAc and sPGM. Initially, we attempted to centrifuge and PBS wash cells, but this resulted in significant cell lysis **(Supplementary table 2)**. To avoid lysis, we utilised a high performance anion exchange chromatography method to detect fluorescent substrates that were sampled directly from the culture media under aerobic conditions (**Figure 3a-c and Supplementary figure 4c)**.

**Figure 3.**
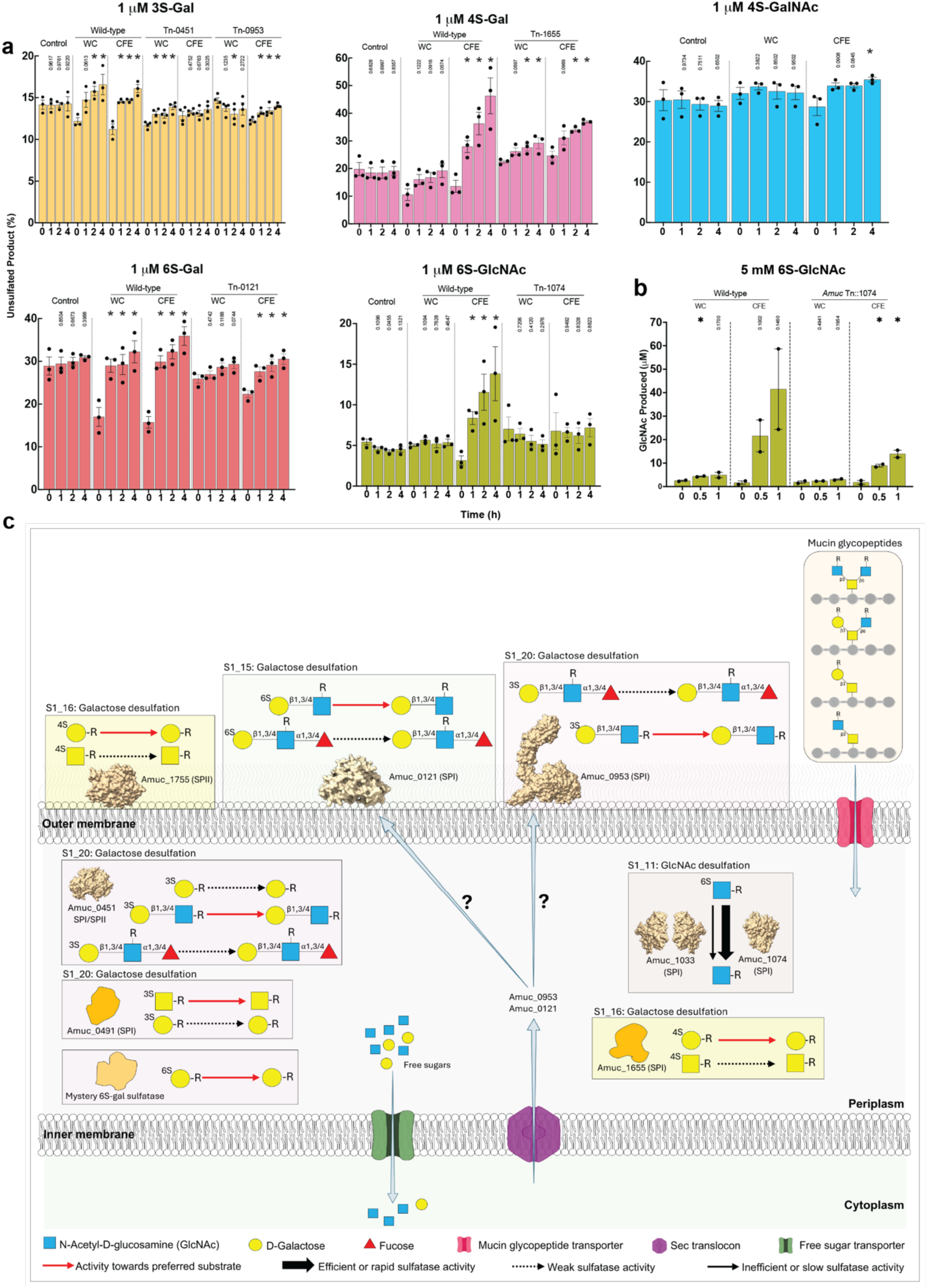
Degradation of 6’S-Lewis antigens and sulfatase cellular location. **a.** HPAEC based cell assays with wildtype and transposon mutant sulfatases. Cells were grown, with sPGM as their carbon source, to mid-exponential phase before being left in aerobic conditions for ∼30 mins to arrest growth. These were either left as whole cells or prepared as CFE and 1 μM of a fluorescent substrates added to the media and sampled periodically. Experiments are at least biological triplicates. Statistical significance of time points versus control was calculated using an unpaired students t-test; an asterix indicates a value <0.5 otherwise the value is stated. **b.** HPAEC based cell assays with wildtype and transposon mutant sulfatases Cells were grown to mid-exponential phase before being left in aerobic conditions for ∼30 mins to arrest growth. These were PBS washed and either left as whole cells or sonicated before adding 5 mM 6S-GlcNAc. These experiments carried out as biological duplicates. Statistical significance of time points versus control was calculated using an unpaired students t-test; an asterix indicates a value <0.05 otherwise the value is stated. **c.** A schematic model of sulfatase cellular location and the substrates targeted.

Assays with wildtype *A. muc* with intact whole cells and CFE indicated that *O*3 desulfation of galactose occurs both extra-and intracellularly **(Figure 3a,c and Supplementary figure 4c)**. Assays with the *amuc0953::Tn* mutant reveal loss of extracellular activity but retention of activity with CFEs. The reciprocal was observed for the *amuc0451::Tn* mutant. These data indicate that Amuc0953^3S-LacNAc^ is responsible for the extracellular activity and Amuc0451^3S-LacNAc^ for intracellular activity. Although Amuc0953^3S-LacNAc^ is predicted to have a SPI signal peptide suggesting a periplasmic localisation, indicating this protein is secreted from the periplasm, and anchored to the membrane, by an unknown mechanism.

Degradation of 4S-Gal was observed from whole cells which increased in CFE indicating the presence of 4S-Gal sulfatase activity both extra-and intracellularly **(Figure 3a and Supplementary figure 4c)**. The *amuc1655::Tn* mutant retained extracellular activity but the increased activity upon cell lysis was markedly reduced. This indicates that Amuc1655^4S-Gal^ is located in the periplasm and Amuc1755^4S-Gal^, which is predicted to have an SPII signal peptide, is present at the cell surface **(Figure 3a)**. Consistent with the biochemical data, only extremely weak 4S-GalNac activity was detected from CFE.

Desulfation of 6S-Gal was observed for whole cells with this activity being lost with the transposon mutant *amuc0121::Tn* **(Figure 3a)** indicating Amuc0121^6S-Gal/GalNAc^ is localised to the cell surface. The open active site of Amuc0121^6S-Gal/GalNAc^ observed in the crystal structure **(Figure 3a Supplementary figure 4a,b)** would facilitate accommodating more complex glycans found at the cell surface. Interestingly, the cell free extracts of *amuc0121::Tn* mutant retain activity indicating the presence of a second, periplasmic, 6S-Gal sulfatase.

Finally, 6S-GlcNAc activity is exclusively observed from CFEs but this activity was lost with the *amuc1074::Tn* mutant indicating Amuc1074^6S-GlcNac^ is located in the periplasm **(Figure 2e and Supplementary figure 4c)**. No 6S-GlcNAc activity was observed for GlcNAc grown cells indicating high GlcNAc concentrations inhibit Amuc1074^6S-GlcNac^ **(Supplementary figure 4c)**. Assays repeated using 5 mM 6S-GlcNAc, instead of 1 µM, show that CFE from *amuc1074::Tn* mutant are able to release GlcNAc at these higher concentrations **(Figure 3b)**. This suggests that the second 6S-GlcNAc sulfatase, Amuc1033^6S-GlcNAc^ is also located in the periplasm. Overall, these results reveal that *O*3, *O*4, and *O*6 desulfation of Gal occurs both extra-and intracellularly whilst the desulfation of 6S-GlcNAc occurs exclusively in the periplasm.

### Amuc1074 is the main 6S-GlcNAc sulfatase in the periplasm

The two S1_11 enzymes, dimeric Amuc1033^6S-GlcNAc^ and monomeric Amuc1074^6S-GlcNAc^ **(Supplementary figure 5a)**, de-sulfate 6S-GlcNAc-BODIPY with similar catalytic efficiencies **(Figure 2a).** This fluorescent label has a six-atom linker and undergoes mutarotation between α and β anomers. However, when we used 6S-GlcNAc β linked to methylumbelliferone (6S-GlcNAc-MU), which more closely mimics a natural glycosidic linkage, we observed a ∼25 fold lower *k*_cat_/*K*_M_ for Amuc1033^6S-GlcNAc^ **(Figure 4a)**. Although data from cellular assays demonstrated that high GlcNAc concentrations inhibited Amuc1074^6S-GlcNAc^ **(Supplementary figure 4c),** and Amuc1033^6S-GlcNAc^ activity was only observed at high substrate concentrations, **(Figure 3b)**, both enzymes had similar affinities for both substrate and product when 6S-GlcNAc-MU was the substrate **(Figure 4a,b)**. The much lower catalytic efficiency observed for Amuc1033^6S-GlcNAc/GlcNS^ is entirely driven by its ∼50 fold lower *k*_cat_ **(Figure 4a).**

**Figure 4.**
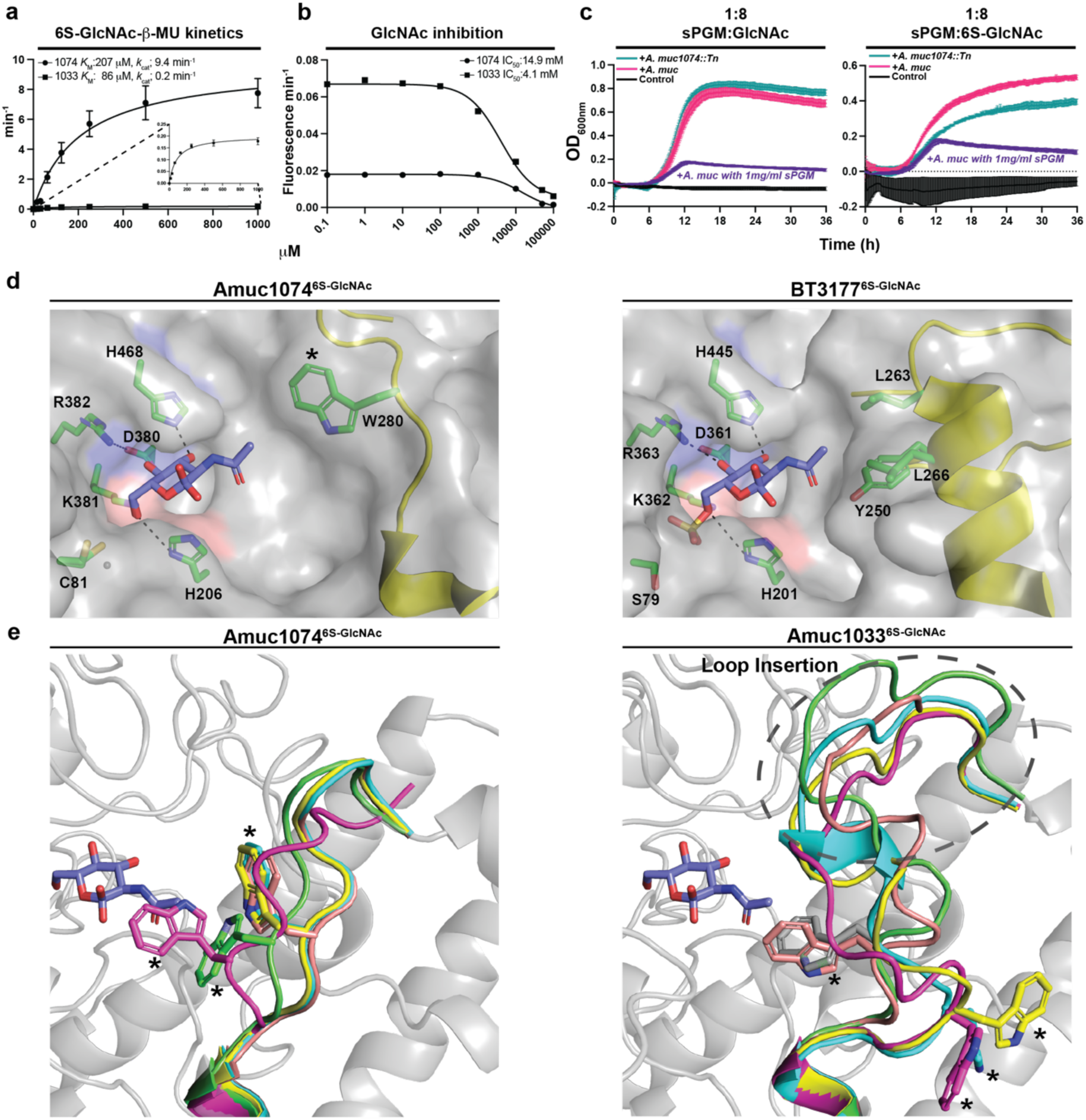
*A. muciniphila* has a low and high activity periplasmic S1_11 sulfatase. **a.** Michaelis-Menten kinetics of Amuc1074^6S-GlcNAc^ and Amuc1033^6S-GlcNAc^ versus 6S-GlcNAc-MU. **b.** IC_50_ curves measuring the ability of GlcNAc to inhibit Amuc1074^6S-GlcNAc^ and Amuc1033^6S-GlcNAc^. **c.** Growth assays of wildtype *A. muc* and Amuc1074::Tn on GlcNAc, 6S-GlcNAc, and 1:8 ratios of sPGM:GlcNAc and sPGM:6S-GlcNAc. The ratios are 1 mg/ml of sPGM and 8 mg/ml of the respective monosaccharide. The growth supported by 1 mg/ml of sPGM is shown for reference. **d.** Crystal structure of the glycan binding site of Amuc1074^6S-GlcNAc^ with GlcNAc compared to the *B. theta* enzyme BT3177^6S-GlcNAc^ (PDB:7P24). A hydrophobic interaction with the *N*-acetyl group is key for high GlcNAc recognition. **e.** Alphafold 2 models of Amuc1074^6S-GlcNAc^ and Amuc1033^6S-GlcNAc^ demonstrating the predicted flexibility of the respective *N*-acetyl recognition loops. An asterix marks the position of the key Trp residue that interacts with the *N*-acetyl group of GlcNAc, which has also been added to the Amuc1074^6S-GlcNAc^ crystal structure in a. All kinetic data are technical triplicates.

Under conditions where GlcNAc is the main carbon source, and in the presence of 1mg/ml sPGM, *amuc1074::Tn* displays no phenotype, but under the same conditions where 6S-GlcNAc is the main carbon source the *amuc1074::Tn* mutant is unable to fully utilise 6S-GlcNAc compared to wildtype with a lower growth rate and maximum O.D. **(Figure 4c)**. This is likely due to the reduced efficiency of Amuc1033^6S-GlcNAc^.

The structure of Amuc1074^6S-GlcNAc^ in complex with GlcNAc reveals it has the same critical recognition features as the *B. theta* enzyme BT3177^6S-GlcNAc/GlcNS^ **(Figure 4d and Supplemental figure 6a)**. The *N*-acetyl recognition loop is highly variable in amino acid composition, and is absent in half of the S1_11 subfamily **(Supplementary figure 7)**, but a hydrophobic character is key for its interaction and Amuc1074^6S-GlcNAc^ and Amuc1033^6S-GlcNAc/^ both deploy a Trp residue to interact with the *N*-acetyl group. The difference in efficiency between the two enzymes is due to Amuc1033^6S-GlcNAc^ containing a loop insertion prior to the *N*-acetyl recognition loop creating greater flexibility, demonstrated by AF2 modelling and molecular dynamics **(Figure 4e and Supplementary figure 6b)**. This leads to a reduced *k*_cat_ through loss of productive energy to the transition state. The presence of Trp in the *N*-acetyl recognition with an extended loop is rare in S1_11 **(Supplementary figure 7)**. *A. muc* has a strong preference for GlcNAc as a carbon source^35^ **(Supplementary figure 8)** and requiring an efficient, periplasmic, 6S-GlcNAc desulfation is in line with this.

### A. muc S1_16 sulfatases have specificity for O4 sulfated D-galactose

Amuc1655^4S-Gal^ and Amuc1755^4S-Gal^ are monomeric (**Supplementary figure 5a)** and have a strong preference for 4S-Gal over 4S-GalNAc **(Figure 2a and Figure 3a)**, in contrast to *B. theta* S1_16 enzymes, which have equal activity on both substrates^31^. The structure of Amuc1755^4S-Gal^ revealed it had the key specificity residues identified in HCM S1_16 enzymes for 4S-Gal/GalNAc recognition **(Figure 5a,b)**. Trp118 stacks against the alpha face of the sugar and Trp462 interacts with *O*3 via the secondary amine of its indole ring. A key difference is the presence of Glu464 which resides on a loop in the motif W_GEX and interacts with *O*2 and limits the accommodation of an *N*-acetyl group thus driving specificity for 4S-Gal **(Figure 5a)**; this residue is also present in Amuc1655^4S-Gal^. Analysis of S1_16 sequence alignments reveal the W_GEX motif is rare and exclusive to the HCM; only 25 out of 800 analysed sequences containing this motif **(Supplementary figure 9)**. Mutation of Glu464 to Ala and Gly enabled the Amuc1755^4S-Gal^ to work on 4S-GalNAc but reduced activity on 4S-Gal **(Figure 5c and Supplementary table 1).** The low activity of the wildtype enzymes on 4S-GalNAc **(Figure 5c)** suggests there is some limited flexibility of the W_GEX loop.

**Figure 5.**
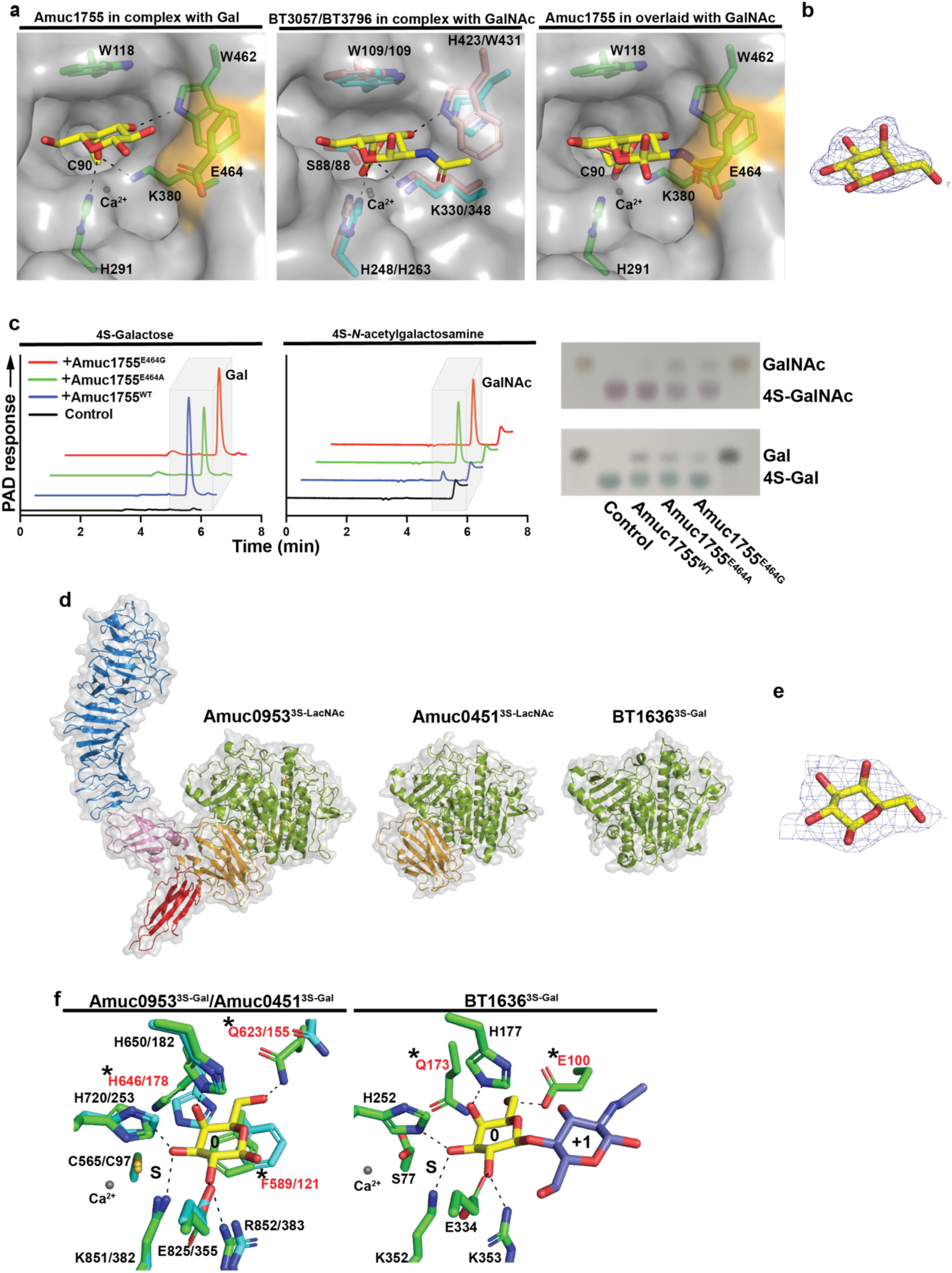
Structural features underpinning specificity in *A. muciniphila* S1_16 and S1_20 sulfatases. **a.** Crystal structure of the glycan binding site of Amuc1755^4S-Gal^ compared with the *B. theta* S1_16 sulfatases that target 4S-Gal and 4S-GalNAc. **b.** 2mFobs-DFc electron density map of D-galactose from the Amuc1755^4S-Gal^ complex contorted to 1σ. **c.** HPAEC (left) and TLC analysis (right) of wildtype and mutant forms of Amuc1755^4S-Gal^ targeting 4S-Gal and 4S-GalNAc. It can be seen E464 is critical for driving 4S-Gal specificity. **d.** Cartoon representations with a transparent surface of A. muciniphila and *B. theta* S1_20 orthologues. **e.** 2mFobs-DFc electron density map of D-galactose from the Amuc0953^3S-Gal^ complex contorted to 1σ. **f.** Stick representation of the glycan binding sites of Amuc0953^3S-LacNac^, Amuc0451^3S-LacNac^, and BT1636^3S-Gal^. The galactose in the left panel is from the Amuc0953^3S-LacNac^ structure and Amuc0451^3S-LacNac^ has been superimposed.

### A. muciniphila multidomain S1_20 sulfatases target 3S-LacNAc

Amuc0451^3S-LacNAc^ and Amuc0953^3S-LacNAc^ are large, monomeric, multidomain sulfatases **(Figure 5d)** that show distinct unfolding stages in their melt curves **(Supplementary figure 5a,b).** To understand the significance of these sizes differences, and specificity, we solved crystal structures of both Amuc0451^3S-LacNAc^ and Amuc0953^3S-LacNAc^ **(Figure 5d,e)**. Amuc0451^3S-LacNAc^ comprises the core S1, α/β/α, fold but with an abutted BACON domain at its C-terminus. Amuc0953^3S-LacNAc^ also has a BACON domain C-terminal to the core sulfatase domain but has two further, smaller, domains one which is an Ig-like domain and the other was unresolved. At the N-terminus Amuc0953^3S-LacNAc^ possesses a large parallel β-helix domain that is connected to the core sulfatase domain via a small β-sheet domain with a single α-helix **(Figure 5d)**. AF3 structure prediction of Amuc0953^3S-LacNAc^ indicates that the final C-terminal domains (CTD) bears resemblance to the CTD recognised by Bacteroidota type IX secretion systems^36^. A comparable system may therefore exist in *A. muc* enabling translocation for Amuc0953^3S-LacNAc^ to the cell surface **(Supplemental Figure 10a,b)**. No 3S-Gal activity could be detected in the media indicating Amuc0953^3S-LacNAc^ is membrane bound at the cell surface **(Supplemental Figure 10c).**

Amuc0451^3S-LacNAc^ and Amuc0953^3S-LacNAc^ have identical binding site sequences and are similar to the critical *B. theta* enzyme BT1636^3S-Gal^ **(Figure 5f)**. His coordinates the Gal axial *O*4, whilst *O*2 is coordinated by Glu and Arg, and *O*6 by a Gln. Amuc0451^3S-LacNAc^ and Amuc0953^3S-LacNAc^ both have a Phe in the glycan binding site that stacks against the Gal sugar ring, replacing Glu100 in BT1636^3S-Gal^. The addition of an interacting Phe residue is accompanied by a Gln to His (replacing Q173 in BT1636^3S-Gal^) covariant mutation, with the His lying perpendicular to Phe, which only occurs in ∼9 % of S1_20 sequences which cluster together and mainly comprise Verrucomicrobia species **(Supplementary figure 11)**. In the Apo Amuc0451^3S-LacNAc^ structure the Phe and His are shifted. The ∼10 fold preference of Amuc0451^3S-LacNAc^ and Amuc0953^3S-LacNAc^ for 3S-LacNAc over 3S-Gal may be due to the flexible stacking/hydrophobic interactions allowing Phe to interact with 3S-LacNAc **(Figure 5f)**.

### The N-terminal domain of Amuc0953 is a new CBM family binding colonic mucin

Parallel β-helix domains are common for CAZymes targeting plant glycans^37^ and GAGs^38^, as well as carbohydrate binding module family (CBM) 89 which binds xylan^39^. However, no enzymatic activity for the N-terminal domain of Amuc0953 could be observed against pectins, xylans, glycosaminoglycans or PGM, nor binding to xylan **(Supplementary figure 12a,b)**. Therefore, we hypothesized that the Amuc0953 N-terminal domain could be the founding member of a CBM family targeting colonic mucin.

To understand the role of the N-terminal parallel β-helix domain we generated two constructs terminating at Pro395 (Amuc0953^P395^) and Pro490 (Amuc0953^P490^) of which Amuc0953^P490^ was successfully crystallised to 2.0 Å **(Figure 6a)**. We conducted pull down assays using PCM and both constructs were pulled down by these colonic mucins, whilst BSA was not **(Figure 6b)**. Furthermore, controls with crystalline cellulose showed the binding was specific to PCM **(Figure 6b)**. Structural comparison to CBM89 reveals Amuc0953^P395^ is not structurally related with calculated R.M.S.D.s of 4.39 and 5.27 Å over 255 and 190 aligned Cα residues, respectively **(Supplementary figure 12c)**. A Blastp^40^ and PSI-Blast^41^ search returned only 206 sequences, of which 201 had a minimum identity of 40 % with a 70 % query coverage **(Figure 6c)**. All sequences were from the genus *Akkermansia* with most from *A. muc* strains suggesting it is a rare sequence largely restricted to *Akkermansia*. Over 90 % of these sequences are associated with a sulfatase domain. This indicates that Amuc0953 N-terminus domain is the founding member of a new CBM family in the *Akkermansia* genus.

**Figure 6.**
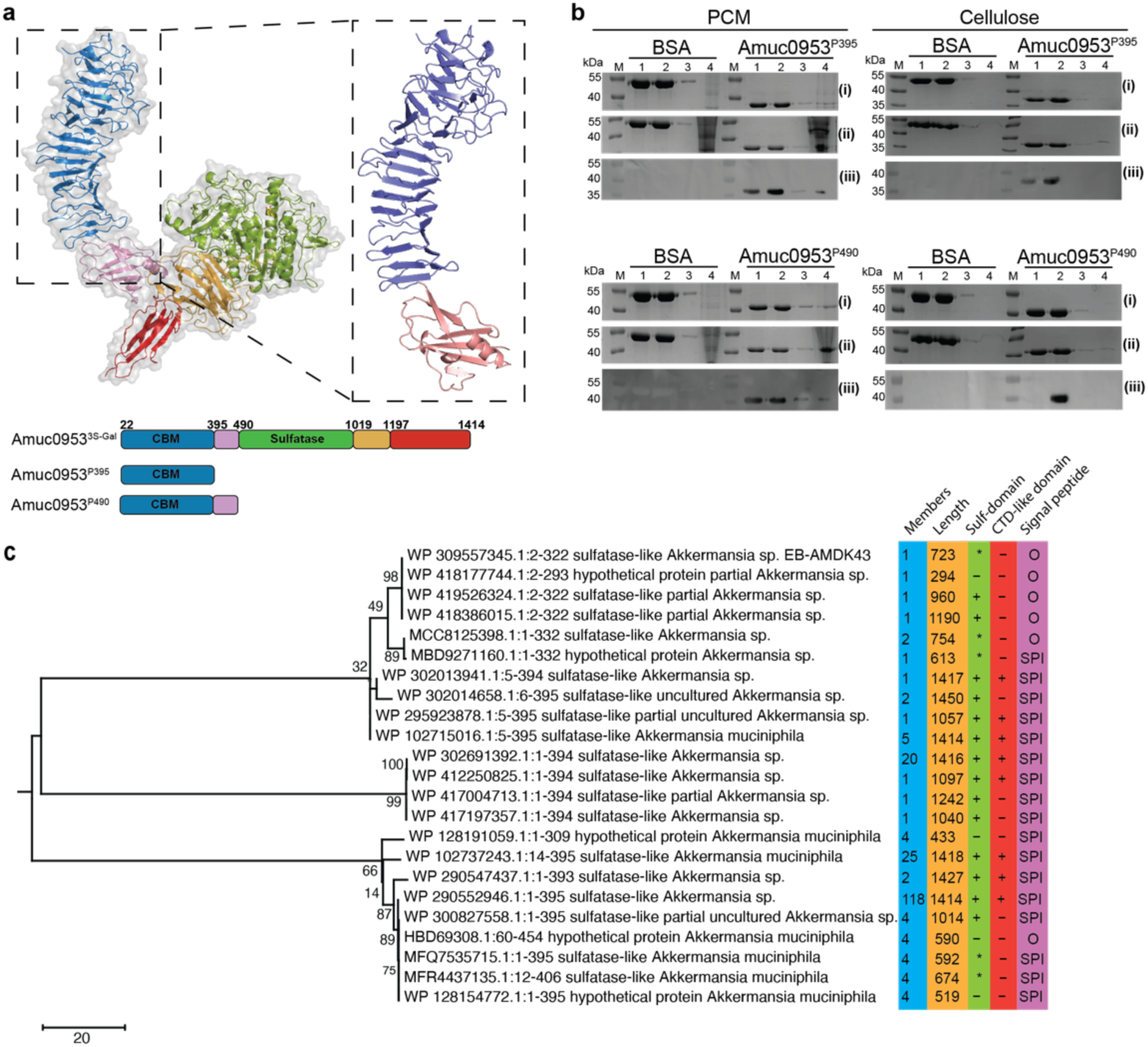
The N-terminus of Amuc0953^3S-Gal^ is a novel carbohydrate binding module. **a.** Cartoon representations with a transparent surface of Amuc0953^3S-Gal^ with the structure of CBMXX, extending to P395, identified in blue and a the linker domain, extending to amino acid P490, shown in salmon. **b.** Pull down assays demonstrating Amuc0953^P395^ and Amuc0953^P490^ bind porcine colonic mucin. (i) SDS-PAGE with 10 μl of pull down sample loaded, (ii) same as (i) but lane 4 has been concentrated ten times, (iii) western blot of (i) using an anti-his antibody. M = marker, 1 = start material: 10 mg/ml glycan plus 10 μM protein, 2 = Supernatant after centrifugation, 3 = supernatant from wash; resuspension of pellet in PBS then centrifugation, 4 = pellet; resuspension of pellet in PBS then boiled. **c.** Unrooted maximum-likelihood tree constructed in MEGA X based on alignment of amino acid sequences from BLAST using the ClusteredNR database. Branch lengths are proportional to the number of substitutions per site. Blue box indicates number of members per clustered sequence and the orange box indicates the sequence length. Green box indicates the presence of the S1_20 sulfatase-like domain in AlphaFold3 models generated from the sequence clusters and the red box corresponds to the presence of a C-terminal domain-like structure (CTD). The purple box indicates if there is a signal peptide present within the sequence as determined by Signal 6.0; SPI indicates translocation through the Sec system to the periplasm whilst ‘O’ indicates a result returned as other. + and – symbols indicate if a domain is present or absent, whilst a * is indicative of the partial/incomplete sulfatase domain. Sequence names and identifiers have been truncated for clarity.

### Growth of carbohydrate sulfatase transposon mutants on mucin substrates

Having determined the catalytic specificities and cellular location of the 8 of the 10 carbohydrate sulfatases we examined the effect of the 8 transposon mutants (*amuc0121*::tn, *amuc0451*::tn, *amuc0565*::tn, *amuc0953*::tn, *amuc1074*::tn, *amuc1182*::tn, *amuc1655*::tn, and *amuc0490-491*::tn) on *A. muc’s* ability to utilise mucin substrates as a sole carbon source. Minor growth, and variable, differences were observed for all mutants grown on GlcNAc and gMOs. **(Supplementary figure 13)**. On sPGM and gpcMOs most mutants displayed no growth differences to wildtype except *amuc0490-491*::tn with had an increased lag phase on both sPGM and gpcMOs, and *amuc1182*::tn which had an increase in lag phase on sPGM. The Tn insertion in *amuc0490-491*::tn spans both genes and the defect observed here is likely linked to *amuc0490* as it is observed across all substrates tested. No activity for Amuc1182^S^^1_19^ could be detected but the phenotype displayed by *amuc1182*::tn suggests the protein has a role in sPGM metabolism not required for gpcMOs **(Supplementary figure 13)**.

## Discussion

*Akkermansia muciniphila* can utilize gMOs and mucins but fails to grow on cMOs. However, recently it has been shown that *A. muc* can grow on trypsin treated colonic mucin suggesting that this bacterium requires the protein backbone to utilise these complex glycans^32^. Our results validate this finding but indicate that *A. muc* requires glycopeptides of a specific size and/or motif to support strong growth on colonic mucin. This indicates that *A. muc* is unable to complete the digestion of this trypsin-treated material to its preferred substrate and may require microbiota driven processing to make them accessible. A recent study showed that *A. muc* transports large mucin portions into its periplasm where they are degraded in a ‘mucinosome’ complex^42^. Our data is consistent with the idea of the transport of glycoproteins/peptides across the membrane and periplasmic degradation where mucin *O*-glycans are broken down to monosaccharides.

By contrast, the *Bacteroides* organism, *B. theta*, grew on all cMO preparations indicating the mucin peptide backbone is superfluous to its transport/degradative machinery. Interestingly these data imply that *A. muc* surface, endo-acting, GH16 enzymes^43,44^, that cleave colonic mucin to produce free *O*-glycans, will produce substrates for *Bacteroides spp.* but not *A. muc* itself. This paints a picture that the colonic mucin substrates of *A. muc* are glycoproteins/peptides with already-processed *O*-glycan chains, and indicates the possibility of mutualistic growth of *A. muc* and other HCM mucin degraders that target free glycan cMOs.

Sulfation of colonic mucin is known to occur on the *O*3, *O*4, and *O*6 of Gal and *O*6 of GlcNAc and presents a steric hindrance to HCM GHs. Although, sulfoglycosidases from GH20 family can cleave 6S-GlcNAc^45^, this sulfated monosaccharide must still be desulfated before for use in central metabolism. De-sulfation of *O*3, *O*4, and *O*6 of Gal occurs at both the cell surface and periplasm in *A. muc*, whilst 6S-GlcNAc de-sulfation is performed exclusively in the periplasm. This surface Gal desulfation could make these residues accessible to HCM galactosidases whilst maintaining 6S-GlcNAc as a bottleneck. This could be beneficial to organisms such as *Rumminococcus torques* that contains no carbohydrate sulfatases and is unable to access sulfated branches of colonic mucin glycans^32^.

In *B. theta* the cell surface S1_20 sulfatase, BT1636^3S-Gal^, is essential for efficient cMO utilisation. In contrast, neither of the two orthologues in *A. muc*, were essential for *A. muc* growth on gpcMOs. Although key enzymes located at the cell surface have been previously described as essential to the utilisation of glycans by *Bacteroides*, it remains unclear if *A. muc* also has such critical enzymes. Both enzymes show a 10 fold preference for the disaccharide 3S-LNB over 3S-Gal which is consistent with data showing *A. muc* cells preferentially bound core 3 structures with LNB epitopes^46^. Additionally, the presence of multiple sulfatases at the *A. muc* cell surface is likely to allow this bacterium to access to different *O*-glycans decorating the imported mucins.

One of the most interesting features of the cell surface Amuc0953^3S-LacNAc^ is the N-terminal domain which comprises a first in class CBM that is exclusively found in the *Akkermansia* genus. These may fulfil a similar role to surface glycan binding proteins observed in *Bacteroides* species for glycan capture^47^, increasing effective substrate concentration for the enzyme and helping Amuc0953^3S-LacNAc^ desulfate terminal 3S-Gal moieties.

The apparent rare specificity for Gal over GalNAc exhibited by the S1_16 enzymes suggests that 4S-Gal, detected in salivary mucins^48^, is a motif commonly encountered by *A. muc*, and the specificity of the *A. muc* sulfatases may give it a competitive edge on this substrate. The S1_11 enzymes in *A. muc* that target 6S-GlcNAc have vastly different activities driven by differences in the flexibility of a key *N*-acetyl loop. Why *A. muc* has an apparent low efficiency 6S-GlcNAc sulfatase is unclear, but it is possible this enzyme targets a larger 6S-GlcNAc containing glycan.

For both gastric and colonic mucin *A. muc* requires a glycoprotein/peptide signature not required by other gram-negative mucin degraders such as *B. theta* indicating a significant difference in acquisition modes. Furthermore, Gal desulfation occurs both outside and inside the cell, potentially making unsulfated Gal more accessible to other HCM species, whilst desulfation of GlcNAc is exclusively periplasmic, and a more selfish pursuit. This is consistent with GlcNAc being a favoured monosaccharide carbon source^35^. Overall, this illustrates significantly different colonic mucin metabolism strategies, involving carbohydrate sulfatases, by *A. muc* and *B. theta* and may begin to shed light on their differing mucin metabolism during symbiosis and dysbiosis.

## Acknowledgements

Funding was provided to AC from the Academy of Medical Sciences/Wellcome Trust through the Springboard Grant (SBF005\1065 163470), the Royal Society through their research grant (RGS\R2\212050), and a Wellcome Trust CDA (225897/Z/22/Z). ASL. was supported by Swedish Research Council Grant (2021-01409) Swedish Society for Medical Research (Svenska Sallskapet for Medicinsk Forskning, Grant S21-0026), Sahlgrenska Academy International Starting Grant (GU2021/1070), and Jeanssons Foundation Grant. ECM, SRS, ASL and CJ were supported by funds from the US National Institutes of Health (R01 DK125445 to ECM). GR was funded by the Estonian Research Council (PUTJD1227). We thank SciLifeLab and BioMS funded by the Swedish research council for providing financial support to the Proteomics Core Facility, Sahlgrenska Academy. S.vd.P was supported by the Swedish Research Council grant (2020-02536) and Sahlgrenska Academy International Starting Grant (GU2021/1070)

## Author contributions

D.D., A.C., N.D.S. and A.S.L. performed protein expression and purification. D.D., A.C., D.P.B., N.D.S., A.S.L, and C.W.E.T conducted quantitative and qualitative enzyme assays.

A.C. performed DSF experiments. A.C., A.S., and D.J.R. performed crystallographic experiments and analysis. A.C. G.S.A.W carried out biophysical characterisation. D.D., A.C.,

L.D. S.R.S., and E.C.M performed bacterial growth experiments. A.C., D.D., S.R.S., and M.O. collected and purified mucin substrates. D.D. carried out cellular localisation experiments. A.S.L., G.R., M.N. and S.vdP. carried out proteomic analysis. A.S.L and C.J. performed mass spectrometry glycomic analyses. M.C. and Z.M. carried out phylogenetic analysis. M.D. and E.A.Y. performed NMR analyses of mucin substrates. A.Case and CB carried out microscopy and cell lysis analysis.

## Data availability statement

The crystal structure datasets generated have been deposited in the in the Protein Data Bank (PDB) under the following accession numbers: 9S7X, 9S85, 9S88, 9S8A, 9SF6, 9S8X, 9S8Y, and 9S8F. Proteomic datasets have been deposited XXXX via PRIDE. Glycomics raw files were uploaded to Glycopost (https://glycopost.glycosmos.org/entry/GPST000622). Information on all other data and materials are contained within the main manuscript and Supplemental Information.

## Code availability statement

No new codes were developed or compiled in this study.

## Methods

### Chemicals and reagents

All sulfated carbohydrates were purchased from Dextra, except O3 sulfated D-galactose that was purchased from Toronto research chemicals.

### Recombinant protein production

Genes were amplified by PCR using the appropriate primers **(Supplementary table 2)** and the amplified DNA cloned in pETite N-His_6_ tags according to the instruction of the Expresso^TM^ T7 cloning and Expression System kit (Lucigen^®^). Recombinant genes were expressed in *Escherichia coli* strains TUNER (DE3) (Novagen), containing the appropriate recombinant plasmid, and cultured to mid-exponential phase in LB supplemented with 50 μg/mL kanamycin at 37 °C and 180 rpm. Cells were then cooled to 16 °C, and recombinant gene expression was induced by the addition of 0.1 or 0.2 mM isopropyl β-D-1-thiogalactopyranoside; cells were cultured for another 16 h at 16 °C and 180 rpm. The cells were then centrifuged at 5,000 × g and resuspended in 20 mM HEPES, pH 7.4, with 500 mM NaCl before being sonicated on ice. Recombinant protein was then purified by immobilized metal ion affinity chromatography using a cobalt-based matrix (Talon, Clontech) and, after a wash with resuspension buffer, eluted with a step gradient of 10-, 50-, and 100-mM imidazole in resuspension buffer. Proteins were then analysed by SDS-PAGE gel for purity and appropriately pure fractions dialysed into 10 mM HEPES pH 7.0 with 150 mM NaCl. Proteins being carried forward for structural experiments were concentrated in centrifugal concentrators with a molecular mass cutoff of 30 kDa, after SDS-PAGE analysis, and loaded onto a 16/60 S200 superdex size exclusion column. Fractions from this were then subject SDS-PAGE analysis and fractions judged to >95 % pure pooled and concentrated in centrifugal concentrators with a molecular mass cutoff of 30 kDa for further downstream structural analyses. Protein concentrations were determined by measuring absorbance at 280 nm using the molar extinction coefficient calculated by ProtParam on the ExPasy server (web.expasy.org/protparam/).

### Site-Directed Mutagenesis

Site-directed mutagenesis was conducted using a modified PCR-based QuickChange protocol. Appropriate primer pairs were designed with the mutation in the middle region and overhangs overlapping the gene sequence of 12 to 20 nucleotides **(Supplementary table 3)**. The plasmid was amplified by PCR using CloneAmp HiFi PCR Premix (Takara Bio) or Phusion^TM^ High-fidelity DNA polymerase (Thermoscientific). Alternatively, mutants were generated using primers that generated blunt ended products with the mutation at the 5’ terminus that produced blunt ended products which were ligated together. A 0.8 % agarose gel was run to visualise the PCR reactions and successful reactions were digested with Dpn I. The digestion reaction (2-5 μL) was transformed into 100 μL of Top10 super chemically competent cells. After recovery, the cells were plated onto LB-agar plates containing kanamycin (50 μg/ml) and grown overnight at 37°C. Colonies were picked, grown up in LB supplemented with 50 μg/ml kanamycin, and the plasmid DNA was isolated and send to sequencing to confirm the correct sequence with the mutation.

### Bacterial growth assays

Cell culture: *Akkermansia muciniphila* ATCC BAA-835was grown in 5ml of Chopped Meat Medium (CMM, **Supplemental table 5**) supplemented with an appropriate carbon source. After recovery, cells were subcultured twice, in CMM with 10 mg/ml *N*-acetyl-D-glucsoamine before setting up an experiment. *Bacteroides thetaiotaomicron* VPI-5482 was recovered in Brain heart infusion media supplemented with haematin. The next day cultures were inoculated in minimal media (MM, **Supplemental table 6**) with an appropriate carbon source for the experiment. Media glycan solutions were made as 2 x stocks and mixed to make 1 x solutions for growth experiments. Cultures were maintained in borosilicate glass test tubes (16×125 mm) at 37℃ inside a Whitley A85 anaerobic workstation filled with a mixture of 10% CO2/10% Hydrogen/Nitrogen. For growth assays, overnight cultures were used, and for whole-cell assays, cells were used at an OD of 0.3-0.4. All growth assays were done in 96-well F-bottom plates, and 300 µL screw top PP vials were used for the whole-cell assays.

### Enzymatic assays analysed by liquid chromatograph with electrospray ionization tandem mass spectrometry (LC-ESI MS/MS)

Assays were performed with 1 μM of recombinant enzymes and 0,5% colonic mucin glycans in 10 mM MES pH 6.5 with 5 mM CaCl_2_ for 16h at 37 °C. The resultant glycans were purified by passing through the graphitized carbon particles (Thermo Scientific) packed on top of a C18 Zip-tip (Millipore). Samples were elutated with 65% acetonitrile in 0.5% trifluoroacetic acid (v/v), dried, and stored at −20 °C until LC-ESI MS/MS analysis. Released glycans were resuspended in 15 μL of Milli-Q water and analyzed by LC-ESI MS/MS using a 10 cm × 250 µm I.D. column, packed with 5 µm porous graphitized carbon particles (Hypercarb, Thermo-Hypersil, Runcorn, UK)). Glycans were eluted using a linear gradient 0–40% acetonitrile in 10 mM NH_4_HCO_3_ over 40 min at a flow rate 6 µl min-1. The eluted O-glycans were detected using an LTQ mass spectrometer (Thermo Scientific, San José, CA) in negative-ion mode with an electrospray voltage of 3.5 kV, capillary voltage of-33.0 V and capillary temperature of 300 °C. Air was used as a sheath gas. Full scan (m/z 380-2000, two microscan, maximum 100 ms, target value of 30,000) was performed, followed by data-dependent MS2 scans (two microscans, maximum 100 ms, target value of 10,000) with normalized collision energy of 35%, isolation window of 2.5 units, activation ρ=0.25 and activation time 30 ms). The threshold for MS2 was set to 300 counts. The data were processed using Xcalibur software (version 2.0.7, Thermo Scientific). Glycans were identified from their MS/MS spectra by manual annotation as previously described^9^. Raw data was uploaded on Glycopost Glycopost (https://glycopost.glycosmos.org/preview/3719534668b934377a9c6, password: 7654). The peak area (the area under the curve, AUC) of each glycan structure was calculated using the Progenesis QI software (Nonlinear Dynamics, Waters Corp., Milford, MA, USA). The AUC of each structure was normalized to the total AUC and expressed as a percentage.

### Size exclusion coupled light scattering determination of molecular weight

Molecular masses were determined using an Agilent Multi-Detector System calibrated with bovine serum albumin. Proteins were separated by size exclusion chromatography using an Agilent BioSEC Advance 300 Å, 4.6 x 300 mm column equilibrated with 20 mM tris(hydroxymethyl)aminomethane-HCl pH 7.4, 150 mM NaCl buffer. Light scattering data was collected at 90° and refractive index used to calculate absolute molecular mass.

### Differential scanning fluorimetry

Thermal shift/stability assays (TSAs) were performed using an Stratagene Mx3005P Real-Time PCR machine (Agilent Technologies) and SYPRO-Orange dye, at a 1:1000 dilution, (emission maximum 570 nm, Invitrogen) with thermal ramping between 20 and 95°C in 0.3°C step intervals every 15 s per data point to induce denaturation of purified, folded, GH139 enzymes and various mutants. The melting temperature (Tm) corresponding to the midpoint for the protein unfolding transition was calculated by fitting the sigmoidal melt curve to the Boltzmann equation using GraphPad Prism, with R^2^ values of ≥0.99, as described in Byrne et al^49^. Data points after the fluorescence intensity maximum were excluded from the fitting. Changes in the unfolding transition temperature compared with the control curve (ΔT_m_) were calculated for each ligand. A positive ΔT_m_ value indicates that the ligand stabilises the protein from thermal denaturation, and confirms binding to the protein. All TSA experiments were conducted using a final protein concentration of 5 μM in 100 mM Bis-Tris-Propane (BTP) or HEPES pH 7.0, and 150 mM NaCl, supplemented with the appropriate ligand concentration. Three independent assays were performed for each protein and protein ligand combination.

### X-ray crystallography experiments

After purification, proteins were carried forward in the same eluent as used for the size exclusion chromatography; 10 mM HEPES pH 7.5, 150 mM NaCl. All proteins were then concentrated in centrifugal concentrators with a molecular mass cutoff of 30 kDa. Sparse matrix screens were set up in 96-well sitting drop TTP Labtech plates (400-nL drops) using an SPT mosquito crystallisation robot, or in sitting drop intelli plates (400-nL drops) using an Arts Robbins gryphon robot. Amuc0451 crystallised in 20 % PEG 3350, 0.1 M Bis-Tris-Propane pH 7.5, and 0.2 M sodium/potassium phosphate at 40 mg/ml with 100 mM D-galactose. Amuc0953 crystallised in 20 % PEG 3350, and 0.2 M sodium iodide at 60 mg/ml D-galactose. Amuc1074 crystallised in the Morpheus screen condition E9, 37.5 % precipitant mix 4, 0.1 M buffer system 2 pH 7.5, and 0.12 ethlene glycols mix at 25 mg/ml with 100 mM N-acetyl-D-glucosamine. Amuc1755 crystallised in 1.6 M sodium citrate at 2.75 mg/ml with 100 mM D-galactose. Amuc0121 at 36 mg/ml with 100 mM D-galactose crystallised in the Morpheus condition H12; 37.5 % precipitant mix 4, 0.1 M buffer system 3 pH 8.5, and 0.1 M amino acids mix. Amuc0121 at 36 mg/ml with 10 mM 6S-Gal crystallised in Morpheus condition D12; 37.5 % precipitant mix 4, 0.1 M buffer system 3 pH 8.5, and 0.1 M alcohols mix. Amuc0121 at 36 mg/ml with 1 mM 6S-LewisA crystallised in Morpheus condition B1; 30 % precipitant mix 1, 0.1 M buffer system 1 pH 6.5, and 0.09 M halogens mix. These crystals were soaked for 24h with 5 mM 6S-LewisA that had been dissolved in the crystallisation condition. Amuc0953^P490^ was crystallised in 2.0 M Ammonium sulfate and 0.1 M Sodium acetate 4.6 at 20 mg/ml; crystals were soaked with 10 mg/ml gpcMOs overnight before fishing, no cryoprotection was used. Crystals from Morpheus conditions do not need additional cryo-protection whilst all other crystals were cryo-cooled with the addition of 20% PEG 400, Ethylene glycol, or glycerol, except Amuc1755 which was cryo-protected with 4.5 M sodium citrate. Data were collected at Diamond Light Source (Oxford) on beamlines I0-3, I0-4, and I24 at 100 K. The data were integrated and scaled with auto processing software at Diamond via Xia2 or autoproc with staraniso pipelines and merged with Aimless^50,51^. Five percent of observations were randomly selected for the R_free_ set. The phase problem was solved by molecular replacement using the program Phaser or Molrep with a model generated through the AF2 colab or the AF3 server. Models then underwent recursive cycles of model building in Coot^52^ and refinement cycles in Refmac5^53^. The models were validated using Coot^52^ and MolProbity^54^. Carbohydrates were made using Jigand^55^. Structural Figures were made using Pymol (The PyMOL Molecular graphics system, Version 2.0 Schrodinger, LLC.) and all other programs used were from the CCP4 suite^56^. The data processing and refinement statistics are reported in **(Supplementary tables 7 and 8)**

### Molecular dynamic simulations

All-atom molecular dynamics simulations were performed in GROMACS 2023 using the Charmm36m forcefield^57^. The explicit solvent systems were set up using CHARMM-GUI^58,59^. Amuc1033 (AlphaFold2^60^), Amuc1074 (PDB ID: 9S8Y) were solvated in water containing 150 mM NaCl and the protonation states of titratable residues were set at pH 7.5. Each system was energy minimised and equilibrated with position restraints at 310.15 K temperature and 1 bar pressure for 5 ns. Systems were then subjected to an unrestrained production run for 100 ns each. The temperature and pressure were maintained by the V-rescale thermostat^61^ and C-rescale barostat^62^. Analysis was performed by in-built Gromacs analysis tools, UCSF Chimera and GraphPad Prism.

### Thin layer chromatography (TLC)

End point assays were analysed by TLC by spotting 2 μL of sample onto silica plates and resolved in butanol:acetic acid:water (2:1:1) solvent system, or formic acid:butanol:water (4:8:1). The plates were dried, and the sugars were visualized using diphenylamine stain (1 ml of 37.5% HCl, 2 ml of aniline, 10 ml of 85% H_3_PO_3_, 100 ml of ethyl acetate and 2 g diphenylamine) and heated at 450°C for 2-5 min with a heat gun.

### Glycan labelling

Sulfated saccharide samples were labelled according to a modification of the method by Das *et al*., reporting the formation of N-glycosyl amines for 4,6-O-benzilidene protected D-gluopyranose monosaccharides with aromatic amines^63^. Briefly, the lyophilised sugar (1 mg) was dissolved in anhydrous methanol (0.50 mL, Sigma-Aldrich) in a 1.5 mL screw-top PTFE microcentrifuge tube and BODIPY-FL hydrazide (4,4-difluoro-5,7-dimethyl-4-bora-3a,4a-diaza-*s*-indacene-3-propionic acid, hydrazide, 0.1 mg, ThermoFischer, λ_ex./em._ 493/503, ε 80,000 M^-1^cm^-1^) was added and the mixture vortexed (1 min), then reacted (65°C, 48 h) in darkness. The products were then cooled and a portion purified by TLC on silica coated aluminium plates (silica gel 60, Sigma-Aldrich, Millipore) resolved with butanol:acetic acid:water (2:1:1). The unreacted BODIPY-FL label (orange on the TLC plate) was identified by reference to a lane containing the starting material (BODIPY-FL hydrazide), allowing differentiation from the putative labelled product (also orange). Products were then separated from unreacted material down a silica column equilibrated with ethyl acetate in with the unlabelled material, but not the labelled sulfated saccharides, run down the column. The labelled product is then eluted off the silica column with methanol and dried (rotary evaporator) to afford the fluorescent, coloured product (bright green in aqueous solution), which was then employed in subsequent experiments.

### Microfluidics de-sulfation assays

Sulfated carbohydrates were labelled at their reducing end with BODIPY-FL which has a maximal emission absorbance of ∼503nm, which can be detected by the EZ Reader via LED-induced fluorescence^49^. Non-radioactive microfluidic mobility shift carbohydrate sulfation assays were optimised in solution with a 12-sipper chip coated with CR8 reagent and a PerkinElmer EZ Reader II system using EDTA-based separation buffer and real-time kinetic evaluation of substrate de-sulfation. Pressure and voltage settings were adjusted manually (1.8 psi, upstream voltage: 2250 V, downstream voltage: 500 V) to afford optimal separation of the sulfated and unsulfated product with a sample (sip) time of 0.2 s, and total assay times appropriate for the experiment. Individual de-sulfation assays were carried out at 28°C and assembled in a 384-well plate in a volume of 80 μl in the presence of substrate concentrations between 0.5 and 20 μM with 100 mM Bis-Tris-Propane, MES, or Tris, depending on the pH required, 150 mM NaCl, 0.02% (v/v) Brij-35 and 5 mM CaCl_2_. The degree of de-sulfation was directly calculated using the EZ Reader software by measuring the sulfated:unsulfated carbohydrate ratio at each time-point. The activity of sulfatase enzymes was quantified in ‘kinetic mode’ by monitoring the amount of unsulfated glycan generated over the assay time, relative to control assay with no enzyme; with sulfate loss limited to ∼20% to prevent of substrate depletion and to ensure assay linearity. *k*_cat_/*K*_M_ values, using the equation *V*_0_LJ=LJ(*k*_cat_/*K*_m_)[E][S], were determined by linear regression analysis with GraphPad Prism software. Substrate concentrations were halved and doubled to assess linearity of the reaction rates to ensure substrate concentrations were significantly <*K*_M_.

### Half maximal inhibitory concentration (IC50) assays

Amuc1033^6S-GlcNAc^ (2 uM) and Amuc1074^6S-GlcNAc^ (50 nM) were assayed against a dilution series of GlcNac (100 nM – 100mM), using 1 uM BODIPY-labelled 6S-GlcNac as a fluorescent substrate. Progress of the reactions were measured by ThermoScientific/Dionex ICS-6000 HPAEC with a Vanquish Fluorescence detector, with separation of sulfated substrate and unsulfated product over a Dionex CarboPac PA-200 3×50mm BioLC guard column at a flow rate of 1 ml min for 3 minutes. A 100 mM sodium hydroxide with 0.2 M sodium acetate mobile phase, with a gradient from 0 - 0.25 min to 1 M sodium acetate which is held until 2 minutes when the column is re-equilibrated back into 100 mM sodium hydroxide with 0.2 M sodium acetate for the remainder of the run was used. Sampling was carried out eight times across a 24hr period (ThermoScientific/Dionex AS-AP autosampler), and relative peak area assessed using Chromeleon 7 (REF). GraphPad prism was used for subsequent data processing and to calculate IC50 values. (REF)

### Linked assays using 6S-GlcNAc **β**-D-methylumbelliferone

Kinetic parameters of Amuc_1033 and Amuc_1074 were monitored via a continuous linked assay using the substrate 4-Methylumbelliferyl-GlcNAc6S (ChemCruz, sc-223640A). Desulfation of the substrate to 4-Methylumbelliferyl-GlcNAc allows it to be hydrolysed by BT_3868^GH^^20^ releasing the fluorophore 4-Methylumbelliferyl (Ex: 355 nm, Em: 460 nm). BT_3868^GH^^20^ mediated hydrolysis is an indirect indicator of desulfation, monitored at 460 nm. A total of 10 substrate concentrations were tested between 0-10 mM in technical triplicate. Enzyme concentrations were 1 μM and 0.1 μM for Amuc_1033 and Amuc_1074 respectively. All assays were carried out in 10 mM HEPES pH 7.5 and 150 mM NaCl. Data acquisition was performed using FLUOstar Omega (BMG LABTECH). Extinction coefficient of 4-Methylumbelliferyl was calculated using a standard curve. 4-Methylumbelliferyl release was plotted as a function of substrate concentration for determination of *K*_M_ and *k*_cat_. Data analysis was performed using GraphPad Prism.

### High performance anion exchange chromatography (HPAEC)

To monitor the liberation sugars from sulfated mono-di-and trisaccharide, the enzyme variants at 5 μM were mixed 1:1 with 5-10 mM of the substrate and incubated overnight at 37 °C. These reaction mixtures were diluted 1:100 in 10 mM Mops 150 mM NaCl, pH 7 and analysed by HPAEC coupled to pulsed amperometric detaction (PAD) for the production of MeFuc. HPAEAC was performed using a Dionex CarboPac PA200 analytical (3 x 250 mm) and guard column (3 x 50 mm) (thermoscientific). Samples were ran for 20 minutes with an isocratic flow of 0.1 M NaOH, before being ran into 100 % 1 M NaOAc for 10 minutes, followed by 5 minutes of 0.5 M NaOH, before a final equilibration step of 0.1 M NaOH for 10 minutes. The flow rate was 0.5 ml/min.

### Cell based enzyme localisation assays via HPAEC with fluorescent detection

Whole-cell assays were performed using *Akkermansia muciniphila* grown in N-acetylglucosamine (BG83, Dextra) and soluble porcine gastric mucin (sPGM) to a final OD of 0.3-0.4. Cells were combined in a 1:1 ratio with 1μM BODIPY-labelled sulfated monosaccharides. For the determination of relative localisation of the sulfatases, both intact cells and cell free extracts were used. Control cells were heat-deactivated at 95℃ for 10 minutes. Sulfatase activity was monitored via BODIPY fluorescence (Ex: 502 nm, Em: 522 nm) through periodic sampling over 4 hours at 37℃. All runs were carried out in a Dionex AS-AP HPLC system (Thermo Scientific) with a Dionex CarboPac^TM^ PA200 BioLC^TM^ 3 x 50 mm Guard column (Thermo Scientific) equilibrated with 100 mM sodium hydroxide (S/4940/17, Fisher Scientific) and 0.2 M sodium acetate (S/2046/50, Fisher Scientific). Sulfated and desulfated monosaccharides were separated using a step gradient of sodium acetate (0.2 M-1 M) over 1.75 ml with a total runtime of 3 minutes. Enzymatic activity was determined by plotting the relative peak area of the desulfated product as a function of time. Monosaccharides tested were 3S-Gal (TRC-G155295, LGC), 4S-Gal (G0012, Dextra), 6S-Gal (G0011, Dextra), 4S-GalNAc (G1054, Dextra) and 6S-GlcNAc (G1004, Dextra). All experiments were carried out in at least three biological replicates. For whole-cell assays to test the activity of Amuc_1033, wild-type and *amuc*1074::Tn mutant cells were exposed to 5 mM of 6S-GlcNAc, and the experiments were conducted in biological duplicates. For all assays, an unpaired two-tailed t-test was utilised to determine statistical significance. Data analysis was performed using GraphPad Prism (v10.5.0).

### Two-colour fluorescence assay for bacterial viability

The LIVE/DEAD™ *Bac*Light™ bacterial viability kit (ThermoFisher Scientific) was used to determine the number of viable and nonviable *A. muciniphila* cells after incubation for 4 hours in chopped meat broth (CMB) or phosphate buffered saline (PBS) pH 7.4. The viability assay was also done on the exponential phase culture from which the *A. muciniphila* were harvested. All bacterial cell containing samples used in the viability assay had approximately the same cell density (OD_600_ ≈ 0.3-0.4). A freshly prepared dye solution containing 10 µM SYTO-9 (dye stains all cells in the population) and 4 µM propidium iodide (dye stains only cells with a comprised membrane) was prepared in either CMB or PBS depending on the experimental condition. Equal volumes (50 µl) of this dye solution and the cell suspension were mixed, then incubated in the dark at room temperature (20°C) for 15 min on a rotary wheel (12 rpm, 80° incline relative to bench top). After staining, the cell suspension was mixed again, and an aliquot (2 µl) of the stained cells was immediately pipetted onto a 1 cm x 1 cm agarose pad and covered with a clean no. 1.5 coverslip (22 mm x 22 mm, Scientific Laboratory Supplies Ltd.). Pads were prepared using low melting point UltraPure agarose (Invitrogen) and un-supplemented M9 medium (48 mM Na_2_HPO_4_, 22 mM KH_2_PO_4_, 8.6 mM NaCl, pH 7.2) according to a published protocol^64^. The cells were imaged by epi-fluorescence microscopy using an Axioskop 40 upright microscope (Zeiss) equipped with a 100x objective (Achroplan NA 1.25, Zeiss), a CoolSNAP EZ camera (Photometrics) and excitation / emission filter sets for the dyes (SYTO-9: Zeiss filter set 10, propidium iodide: Zeiss filter set 43). If the cell density was low in a microscopy sample, the corresponding stained cell suspension was centrifuged (5000 x *g* for 5 min at 20°C) and a portion of the supernatant (∼90 µl) was carefully removed. The cell pellet was resuspended in the remaining supernatant (∼10 µl) and a fresh agarose pad mounted sample was prepared for microscopy as above. Image stacks (35 ms exposure, 100 frames, no interval delay) were collected using MicroManager (version 1.4.16)^65^. The image stacks were averaged in FIJI (version 2.14)^66^ and the number of stained cells in each fluorescence channel was manually counted. The calculated number of viable cells for each condition was equal to the number of SYTO-9-stained cells minus the number of propidium iodine-stained cells. Three biological replicates (n = 3) were analysed for each experimental condition.

### Colonic mucin extraction

Frozen porcine colons stretching ∼ 1 m in length from the anus were thawed, cut open, and washed with tap water before having the mucus scraped using the back edge of a blunt instrument. The scraped material was then either frozen or carried forward into an extraction buffer, containing 6 M GuHCl, 5 mM EDTA, and 0.01 M sodium dihydrogen phosphate at pH 6.5 that was 5-10 times the in volume versus the mass of material, and homogenised with a blunt instrument. The material was stirred slowly for 1-2 h at room temperature before centrifugation at 15,000 x g for 30 mins, supernatant discarded and fresh, cold, extraction buffer added to the insoluble pellet. This process is repeated at least 5 times, or until the last two extractions are clear. The extracted material is then solubilised in 0.5 L of reduction buffer, containing 6 M GuHCl, 5 mM EDTA, and 0.1 M Tris at pH 8.0 with 25 mM DTT left of ∼ 5 hours at 37°C. To this 62.5 mM of iodoacetamide is added and the mixture stirred slowly at room temperature in the dark overnight. The material is then centrifuged at 10,000 x g for 30 mins at 4°C and the insoluble material discarded. The supernatant containing solubilised mucins were extensively dialysed against MQ water at least 6 times before being lyophilised and a weight calculated.

### Release of *O*-glycans by **β**-elimination

Mucin (25 g) was added to 1L of 100 mM Tris pH 7.0 and the mixture was autoclaved and allowed to cool to 65°C before 100 mg of Proteinase K was added and incubated overnight at 65°C. Insoluble material was removed by centrifugation at 10,000 x g for 30 minutes before adding 1 M NaBH4 and 0.4 M NaOH and left overnight at 65°C. The mixture is then neutralised to pH 7.0 using HCl before centrifugation at 10,000 x g for 30 minutes and filtering through a 0.2 micron filter. The material was extensively dialysed (1 kDa cutoff) against water before being lyophilised and a weight calculated.

### NMR analysis

NMR spectra were recorded using a Bruker NEO 700 MHz spectrometer, or a Bruker Avance II+ or 800 MHz Avance Neo spectrometer. Both were fitted with a 5 mm TCI nitrogen cryoprobes. In all cases, a sample of the mucin samples in 600uL of D_2_O was prepared. Data was recorded at 298k, and samples were stored at 4 °C when not actively being measured. For experiments on the Bruker NEO 700 MHz spectrometer: 1D proton spectra were acquired with 16 scans, both with and without presaturation solvent suppression (Bruker sequence ZGPR). 2D 1H-13C HSQC were acquired (Bruker sequence hsqcedetgpsisp2.4). For SPGM spectra were collected with 8 scans and 1024 increments in the indirect dimension. In the case of SPCM and GMO 32 scans and 512 increments were used. For experiments on the Bruker Avance II+ or 800 MHz Avance Neo spectrometer: 1D proton spectra and 2-D HSQC NMR experiments were performed on fractions solubilised in 700 uL D2O (5-mm NMR tubes). Chemical shifts are quoted relative to DSS (0 ppm) using zpgr (1-D 1H) and hsqcetgpsisp (HSQC) pulse programs. Spectra were processed using Bruker TopSpin 3.4 and 4.0 software.

### Glycoprotein gel staining

Glycoprotein gel staining was performed using the Pro-Q^®^ Emerald 300 Glycoprotein Gel and Blot Stain Kit (Molecular Probes, P21857). Samples were stained according to the given protocol. Briefly, mucin and mucin oligosaccharides were separated in a 12.5% SDS-PAGE gel. Post-run, the gel was subjected to fixation and a wash. The gel was then exposed to an oxidising solution, followed by 3 washes. Finally, the gel was exposed to the staining buffer for 90 minutes, followed by 3 additional washes and imaged using a 300 nm UV transilluminator.

### Isothermal titration calorimetry

To quantify the binding affinity of Amuc0953^P395^ to arabinoxylan (5 mg/ml) polysaccharide ITC was performed using a MicroCal Auto-iTC200 (Malvern). The protein sample (50 μM) was stirred at 750 rpm and 400 μL of sample was added to the loading plate, alongside 120 μL of titrant. An initial injection (0.5 μL) of titrant was added prior to an additional 19 injections (2 μL). Titrations were carried out in 50 mM Tris-HCl buffer, pH 8.0, at 25LJ°C. Titrant to buffer, buffer to protein and buffer to buffer control experiments were performed. Data analysis was performed using MicroCal PEAQ-ITC Analysis software (Malvern).

### Pull down binding assays

Protein, at a concentration of 10 μM, was mixed with a polysaccharide at 10 mg/ml and incubated together for 30 minutes in 10 mM HEPES pH 7.0 and 150 mM NaCl. Samples were then centrifuged at 17,000 x *g*, supernatant removed, and the pellet washed and resuspended in 1 ml of buffer. Samples were then centrifuged at 17,000 x *g* again supernatant removed, and the pellet resuspended in either 1 ml or 0.1 ml of buffer as needed. A 10 μl sample was taken at each stage, and these ran on a 12.5 % Tris-glycine SDS-PAGE gel and developed with Coomassie blue stain. Samples were also subjected to western blotting by being transferred to a nitrocellulose membrane, 0.45 μM. Membranes were initially blocked with 5 % milk powder suspended in PBS for 1 h before being washed 3 times with PBS and 0.1 % tween for 5 minutes each. Membranes were then blotted with a mouse anti-his antibody (ThermoFisher MA1-21315), 1:5000 dilution, in PBS with 5 % milk powder overnight at 4°C. Membranes were the washed 3 times with PBS and 0.1 % tween for 5 minutes each and blotted with rabbit anti mouse conjugated to HRP (ThermoFisher 31450), and developed with Clarity™ Western ECL Substrate (BioRad 1705060).

### Phylogenetic and sequence analyses

Phylogenetic analyses were performed as in^9^. In summary, we selected a representative number of sequences within each subfamily to avoid identical sequences still covering the taxonomic diversity: a) for S1_11, 955 sequences were selected from the SulfAtlas database and 411 positions were used for phylogeny; b) for S1_16, 800 sequences were selected from SulfAtlas and 342 positions were used for phylogeny; and c) for S1_20, 848 sequences present in SulfAtlas were selected, and 364 positions were used for phylogeny. In each case, the sequences were aligned by MAFFT v.7^67^ using L-INS-i algorithm. The multiple sequence alignments were visualized by Jalview software v.11.0^68^, non-aligned regions were removed, and the above listed respective numbers of positions were used for the phylogeny. Phylogeny was made using RAxML v. 8.2.4^69^. The phylogenetic trees were built as described in^9^. Subsequently, the aligned sequences were analysed using Jalview and figures created with FigTree v1.4.4^70^.

### Proteomics analyses

The microbial cell pellets were dissolved in 100 uL of lysis buffer (2% SDS (sodium dodecyl sulfate), 5 mM EDTA (ethylenediaminetetraacetic acid), 0.2 M HEPES (4-(2-hydroxyethyl)-1-piperazineethanesulfonic acid), pH 8.0) and shaken at 2000 rpm at room temperature for 5 min, followed by ultrasonication for 4 min (30 sec on/off pulses, 40% amplitude, Fisherbrand Model 505, Thermo Fisher Scientific). Cell lysates were quantified with BCA (bicinchoninic acid) assay (Pierce^TM^ BCA Protein Assay Kit). 25 ug of protein was transferred to a low-binding tube and volume was normalized to 100 μL with lysis buffer. Proteins were reduced and alkylated by the addition of 5 mM TCEP (tris(2-carboxyethyl)phosphine) and 20 mM IAA (iodoacetamide) followed by incubation at 60 °C for 45 min (covered from light). The proteins were digested with trypsin and LysC (Pierce^TM^ Trypsin/Lys-C Protease Mix, MS Grade) on magnetic beads (1:1 mix of Sera-Mag SpeedBead carboxylate-modified [E3] and [E7] magnetic particles, Cytiva) according to single-pot, solid-phase-enhanced sample-preparation (SP3) protocol^71^. Peptide yield was measured with a microvolume spectrophotometer (Nano Drop 2000; Thermo Fisher Scientific) at 280 nm wavelength. The samples were acidified with TFA (trifluoroacetic acid) to final concentration of 0.5% and 15 μg of peptides were cleaned and stored on C18-StageTip filters at-20 °C prior to analysis^72^. Peptides were analysed by LC/MS-MS with an EASY-nLC 1200 system (Thermo Fisher Scientific) coupled to a Q-Exactive HF-X mass-spectrometer (Thermo Fisher Scientific) through a nanoelectrospray ion source. In-house column (200 mm × 0.075 mm inner diameter) packed with Reprosil-Pur C18-AQ 3 μm particles (Dr. Maisch) was used for peptides separation, using the following gradient: 5-30% B over 120 min, 30-45% B over 15 min, 45-100% B over 5 minutes and held for additional 10 min on 100% of B, at flow rate of 250 nl/min (A: 0.1% FA, B: 80% ACN, 0.1% FA). The mass spectrometer was operated at 250 °C capillary temperature and 2.0 kV spray voltage. Full mass spectra were acquired over a mass range from *m/z* 400 to 1600 with resolution of 60 000 (*m/z* 200) after accumulation of ions to a 3e6 target value based on predictive AGC from the previous full scan. Twelve most intense peaks with a charge state ≥2 were fragmented in the HCD collision cell (normalized collision energy of 27%), and tandem mass spectrum was acquired with a resolution of 15 000, AGC target value 1e5. Dynamic exclusion was set to 30 s and the maximum allowed ion accumulation times were 20 ms for full MS scans and 50 ms for tandem mass spectra. Protein sequences for *Akkermansia muciniphila* ATCC BAA-835 were downloaded from the NCBI GenBank database (downloaded 2025.04.21) and supplemented with an in-house database containing human and mouse mucin sequences (https://www.medkem.gu.se/mucinbiology/databases/)^73^. The MS/MS spectra were analysed with MaxQuant (v2.6.5.0)^74^. Searches were performed using trypsin as an enzyme and maximum 2 missed cleavages. Precursor tolerance of 20 ppm was used in the first search for recalibration, followed by 4.5 ppm for the main search and 20 ppm for fragment ions. Carbamidomethylation of cysteine was set as a fixed modification and methionine oxidation and protein N-terminal acetylation were set as variable modifications. The required false discovery rate (FDR) was set to 1% at both peptide and protein levels and the minimum required peptide length was set to seven amino acids. The protein identification and quantification data was further analyzed with Perseus (version 2.0.11)^75^. The same mass spectrometry method as described above, with the exception of a shortened 40-min gradient, was used to assess the presence of glycans and glycopeptides in the porcine colonic glycoprotein fraction. Mass spectrometry data were converted to peaklists using MSConvert and spectra containing following oxonium or immonium fragment ions: HexNAx 138.055, proline 70.065 and threonine 74.006 +/-0.001 Da were extracted with a custom python script. The mass spectrometry proteomics data have been deposited to the ProteomeXchange Consortium via the PRIDE partner repository^76^ with the dataset identifier PXDXXX.

**Supplementary figure 1. Growth profiles and protein expression of *A. muciniphila* on variable mucin structures a.** Growth profiles of *A. muciniphila* on low sulfation and high sulfation soluble mucins. All substrate concentrations are 5 mg/ml unless stated. **b.** HSQC 2-D NMR profiles of various mucin or mucin derived samples examining their protein to glycan content. **c.** Proteomic analysis of the differential regulation of genes compared to growth on monosaccharide and mucin derived substrates.

**Supplementary figure 2. The pH optima of *A. muciniphila* carbohydrate sulfatases.** Graphs showing the pH optima for *A. muciniphila* sulfatases. All reactions were performed using 100 mM of the appropriate buffer supplemented with 150 mM NaCl and 5 mM CaCl_2_. A substrate concentration of 1 µM was used and an enzyme concentrations of between 0.1 – 2 µM was deployed depending on the enzyme.

**Supplementary figure 3. Crystal structures and activity of Amuc0121^6S-Gal^ and cellular localisation assays of *N*-acetyl-D-glucosamine grown *A. muciniphila* cells.**

**a, b.** Surface representation of the Amuc0121^6S-Gal^ crystal structure in complex with D-galactose and *O*6 sulfated D-galactose, respectively. Key residues are shown as sticks beneath. In orange is the galactose recognition triad whilst green highlights the hydrophobic patch that accommodate Fuc (L-fucose); 6S-Gal is O6 sulfated D-galactose whilst GlcNAc is *N*-acetyl-D-glucosamine. In each case the the weighted 2mFobs-DFc electron density map, contoured at 1 σ, is shown on the right. **c.** HPAEC based cell assays with wildtype and transposon mutant sulfatases. Cells were grown with, *N*-acetyl-D-glucosamine as their carbon source, to mid-exponential phase before being left in aerobic conditions for ∼30 mins to arrest growth. These were either left as whole cells or sonicated and 1 μM of a fluorescent substrates added to the media and sampled periodically. Experiments are biological triplicates. Statistical significance of time points versus control was calculated using an unpaired students t-test; an asterix indicates a value <0.5 otherwise the value is stated.

**Supplementary figure 4. Activity of *A. muciniphila* sulfatases against model sulfated oligosaccharides.**

High performance anion exchange chromatography traces showing the activity of S1 *A. muciniphila* sulfatases on model substrates representing sulfated linkages found in colonic mucin. HPAEC traces are from qualitative overnight assays using 5 mM substrate and 5 μM enzyme carried out a 37°C in 5 mM MOPS pH 7.0 with 150 mM NaCl and 5 mM CaCl_2_.

**Supplementary figure 5. Size determination and biophysical properties of *A. muciniphila* S1 sulfatases**

**a.** Size-exclusion chromatography coupled to light scattering profiles for *A. muciniphila* S1 sulfatases. In each case 5 mg/ml protein was utilised. **b.** Differential scanning fluorimetry profiles for Amuc0953^3S-LacNAc^ and Amuc0953^3S-LacNAc^ showing multiple, distinct unfolding events.

**Supplementary figure 6. Phylogenetics, structure and dynamics of the S1_11 enzymes *N*-acetyl recognition loop structure.**

**a.** Cartoon and stick representation of the N-acetyl recognition loop of Amuc1074^6S-GlcNAc^ in complex with GlcNAc; the weighted 2mFobs-DFc map has been contoured at 1 σ. **b.** Molecular dynamic simulations of unliganded Amuc1033^6S-GlcNAc^ and Amuc1074^6S-GlcNAc^ showing the R.M.S.F. values per residue with a zoomed in graph of the N-acetyl recognition loop.

**Supplementary figure 7. Phylogenetic tree of the S1_11 subfamily**

**a.** Phylogenetic tree of 970 selected, most representative, sequences belonging to S1_11. The colours distinguish the sequences according to different lengths of the *N*-acetyl recognition loop, which in addition is highly variable in amino acid composition. Absence of loops and those with less than 5 amino acids are coloured in purple, those having 5 to 11 amino acids are coloured in green. Sequences having loops with lengths ranging from 12 to 19 and hydrophobic patches similar to Amuc1074 are coloured in yellow and those having loops longer than 19 amino acids (in general 20 to 25) are coloured in orange. Since the hydrophobic character (Trp in Amuc1074 and Amuc1033) is key for its interaction with the *N*-acetyl group of GlcNAc, some isolated sequences having loops without hydrophobic character are highlighted by pink lines.

**Supplementary figure 8. Growth of *A. muciniphila* on monosaccharide growth substrates.**

**a.** Bar chart showing the maximum OD_600nm_ each of the tested monosaccharides can allow *A. muciniphila* to reach **b.** Individual growth curves of *A. muciniphila* grown on monosaccharide substrates. All monosaccharides are at 10 mg/ml and the data are technical triplicates.

**Supplementary figure 9. Phylogenetic tree of the S1_16 subfamily**

Phylogenetic tree of 815 selected, most representative, sequences belonging to S1_16. Human and animal microbiome associated bacteria are coloured in light purple (mainly gut, faeces, and oral cavity), the dark purple are exclusively gut bacteria and are those sequences carrying the W_GEX stretch containing E464; light blue are aqueous environmental bacteria (i.e. waste water, rivulet, mangrove, Chinese sea sediment, etc). Notably, three sequences among these aquatic environments posses the W_GEX pattern (purple dots). All other sequences, coloured in yellow are bacteria isolated from various environmental, mainly soil samples.

**Supplementary figure 10. Comparison of the Amuc0953^3S-LacNAc^ crystal structure with its AF3 model, and 3S-Gal activity in the media. a.** Overlay of the crystal structure of Amuc0953^3S-LacNAc^ and its AF3 predicted model **b.** Comparison of the C-terminal domain (CTD) of Amuc0953^3S-LacNAc^ with the CTD from type IX secretion systems. **c.** Filtered media from *A. muciniphila* cells grown on soluble porcine gastric mucin to mid exponential was tested for its ability to desulfate 3S-Gal.

**Supplementary figure 11. Phylogenetic tree of the S1_20 subfamily**

Phylogenetic tree of 848 selected, most representative, sequences belonging to S1_20. The sequences carrying the concomitant Gln/His & Glu/Phe mutations are highlighted. Three sequences from Streptomyces sp. among the neighbouring group, all belonging to the Actinomycetota phylum, also contain the Gln/His & Glu/Phe mutations and are marked by blue lines and dots. The sequences of the branch highlighted in blue all belong either to the Bacteroidota, Planctomycetota or Verrucomicrobiota phylum.

**Supplementary figure 12. Testing of Amuc0953^P395^ for catalytic and bidnding activity against selected host glycans, and comparison to CBM89.**

**a.** Thin layer chromatography of various polysaccharides incubated with Amuc0953^P395^. For highly charged and sulfated glycans a second solvent system was used to ensure that any oligosaccharide products produced had the ability to migrate. All reactions were in 10 mM HEPES pH 7.0, with 150 mM NaCl, and 10 μM Amuc0953^P395^. **b.** Isothermal titration calorimetry of 50 μM Amuc0953^P395^ against 5 mg/ml wheat arabinoxylan. **c.** structural overlay of Amuc0953^P395^ with CBM89 (PBMDCECB_09513; PDB:7JIV), the only other CBM to have a β-helix fold. **d.** Comparison of the area housing the key binding residues in Amuc0953^P395^ and CBM89 (PBMDCECB_09513) from the JCE algorithm overlay.

**Supplementary figure 13. Growth of transposon mutants on mucin substrates.** Bacterial growth curves of *Akkermansia muciniphila* ATCC BAA-835 and sulfatase transposon mutants on gastric and colonic mucin derived substrates. All growth curves are technical triplicates performed in CMM as a base media supplemented with 5 mg ml^-1^ of the appropriate carbon source as listed. GlcNAc = *N*-acetyl D-glucosamine, gMOs = gastric mucin oligosaccharides, sPGM = soluble porcine gastric mucin, and gpcMOs = glycoprotein porcine colonic mucin oligosaccharides.

