## Supplemental information for "The role of *Akkermansia muciniphila* sulfatases in colonic mucin utilisation"

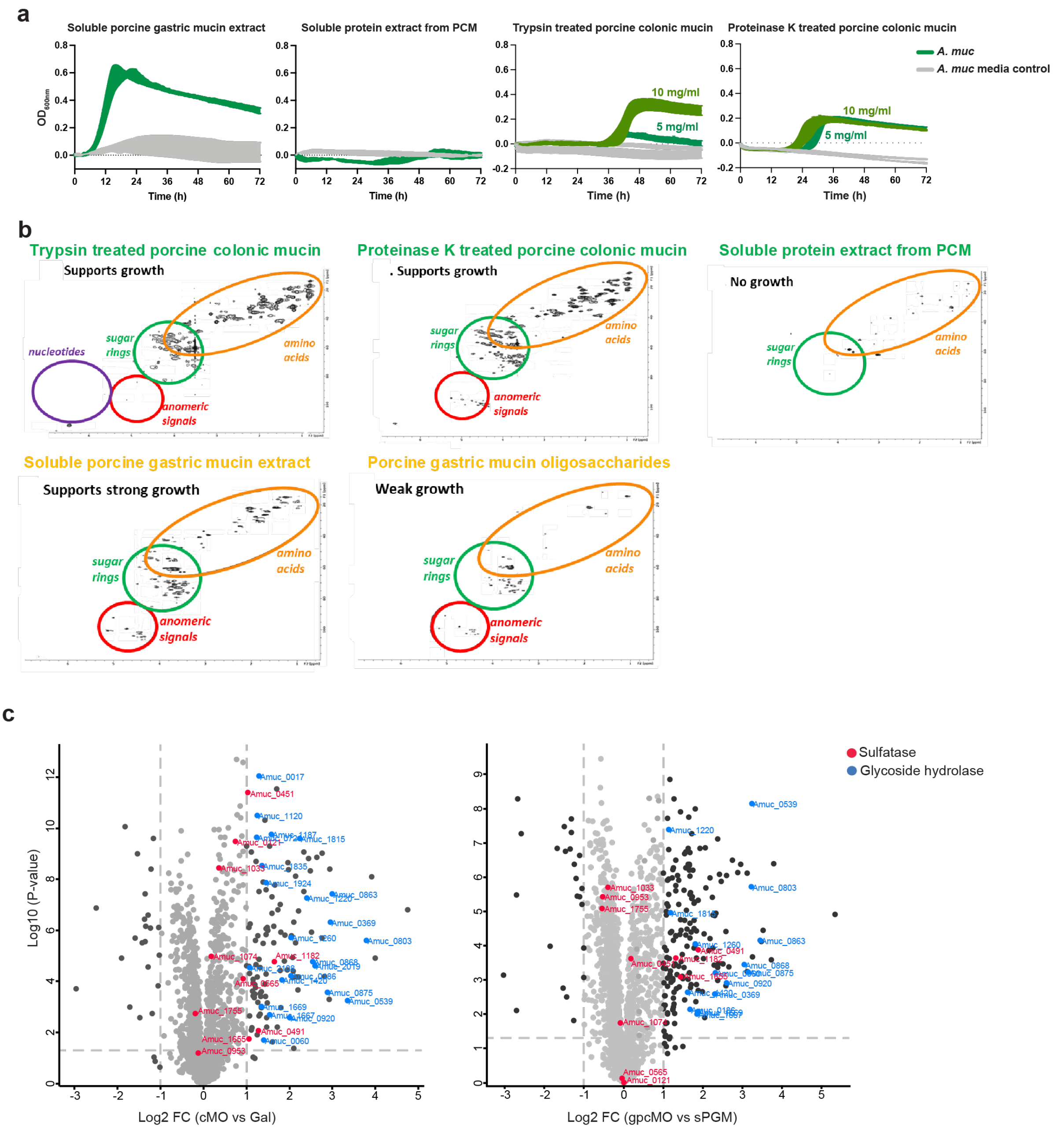


**Supplementary figure 1. Growth profiles and protein expression of *A. muciniphila* on variable mucin structures**

**a.** Growth profiles of *A. muciniphila* on low sulfation and high sulfation soluble mucins. All substrate concentrations are 5 mg/ml unless stated. **b.** HSQC 2-D NMR profiles of various mucin or mucin derived samples examining their protein to glycan content. **c.** Proteomic analysis of the differential regulation of genes compared to growth on monosaccharide and mucin derived substrates.

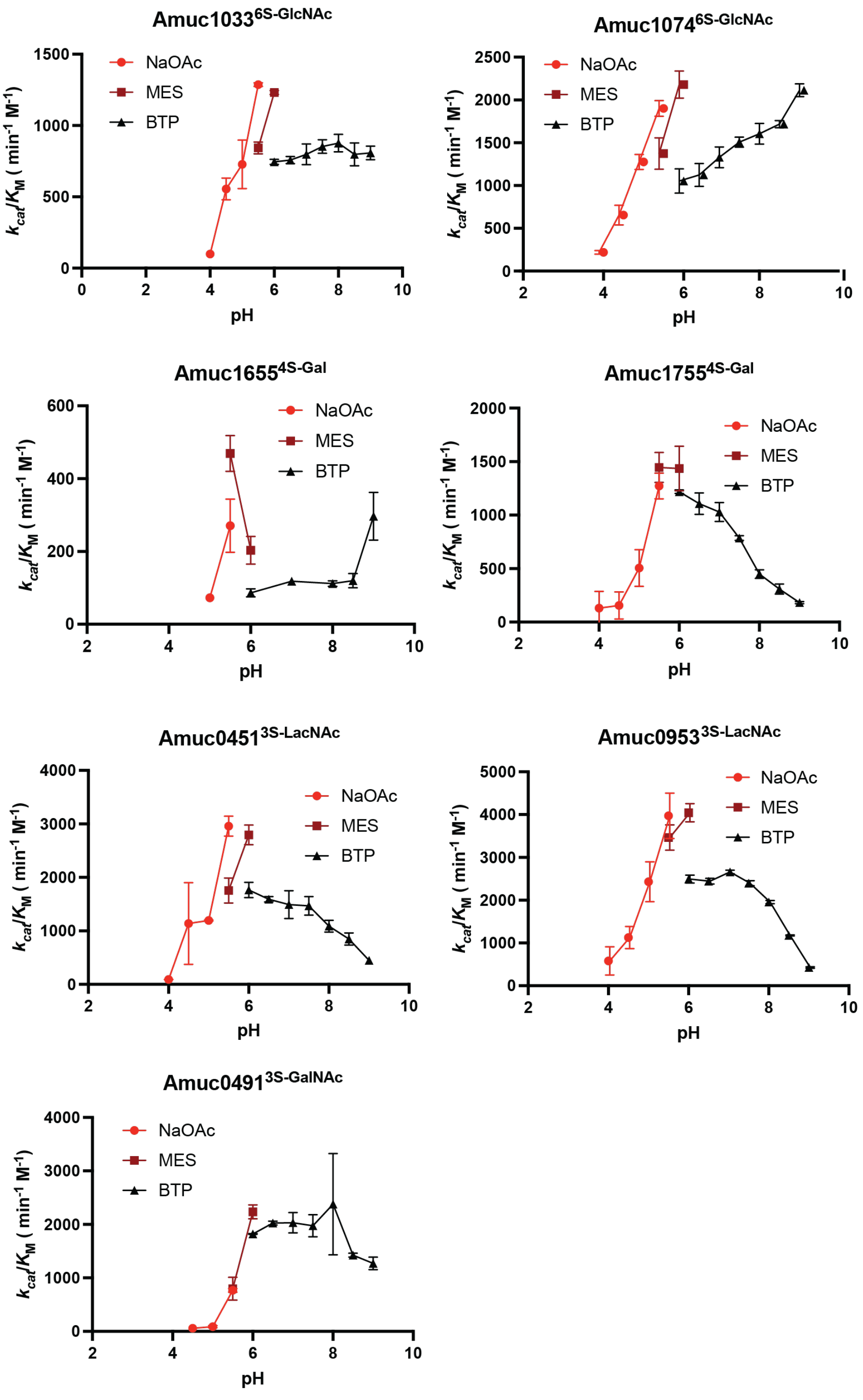


**Supplementary figure 2. The pH optima of *A. muciniphila* carbohydrate sulfatases.** Graphs showing the pH optima for *A. muciniphila* sulfatases. All reactions were performed using 100 mM of the appropriate buffer supplemented with 150 mM NaCl and 5 mM $CaCl_2$. A substrate concentration of 1 µM was used and an enzyme concentrations of between 0.1 – 2 µM was deployed depending on the enzyme.

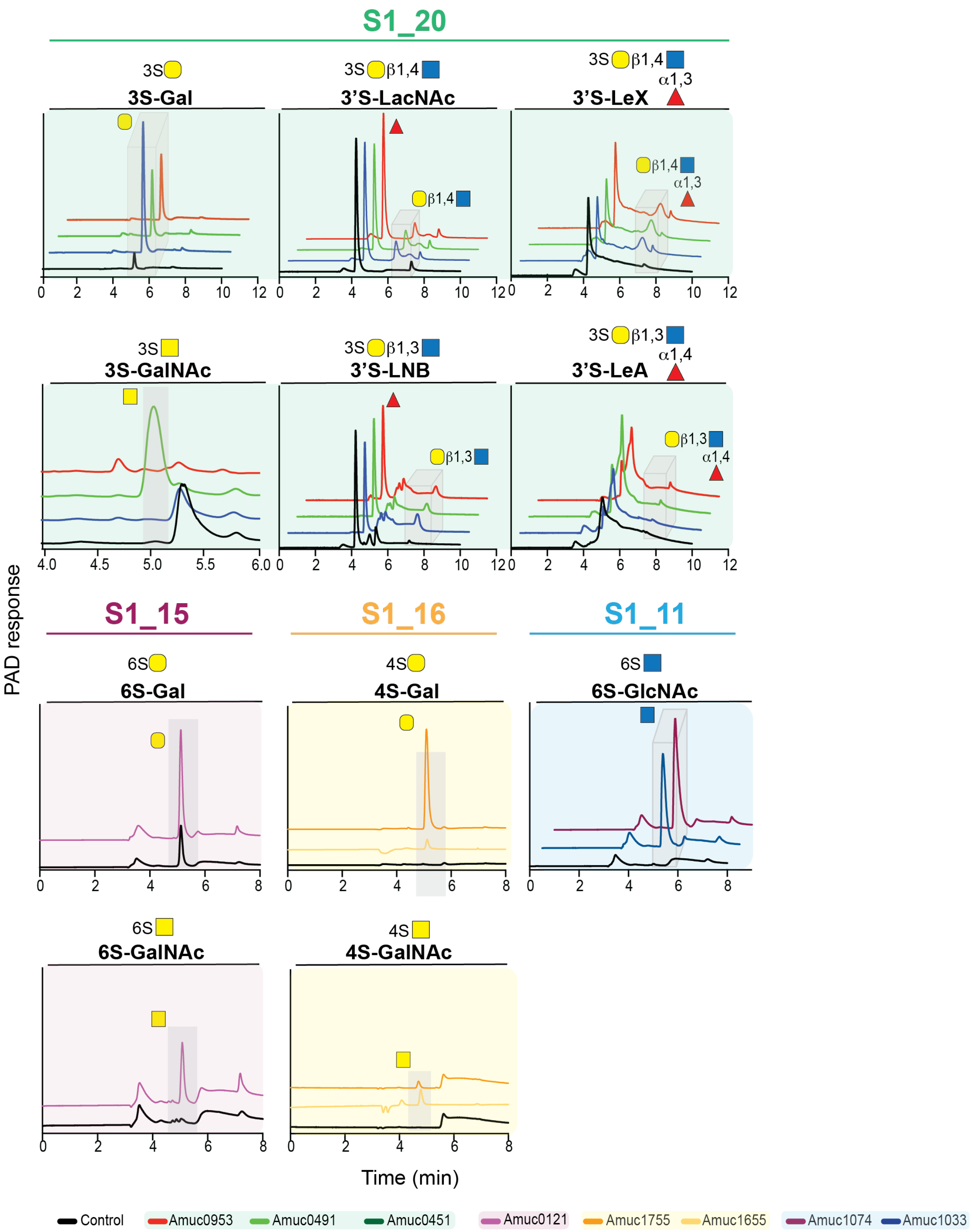


**Supplementary figure 3. Activity of *A. muciniphila* sulfatases against model sulfated oligosaccharides.**

High performance anion exchange chromatography traces showing the activity of S1 *A. muciniphila* sulfatases on model substrates representing sulfated linkages found in colonic mucin. HPAEC traces are from qualitative overnight assays using 5 mM substrate and 5 μM enzyme carried out a 37ºC in 5 mM MOPS pH 7.0 with 150 mM NaCl and 5 mM $CaCl_2$.

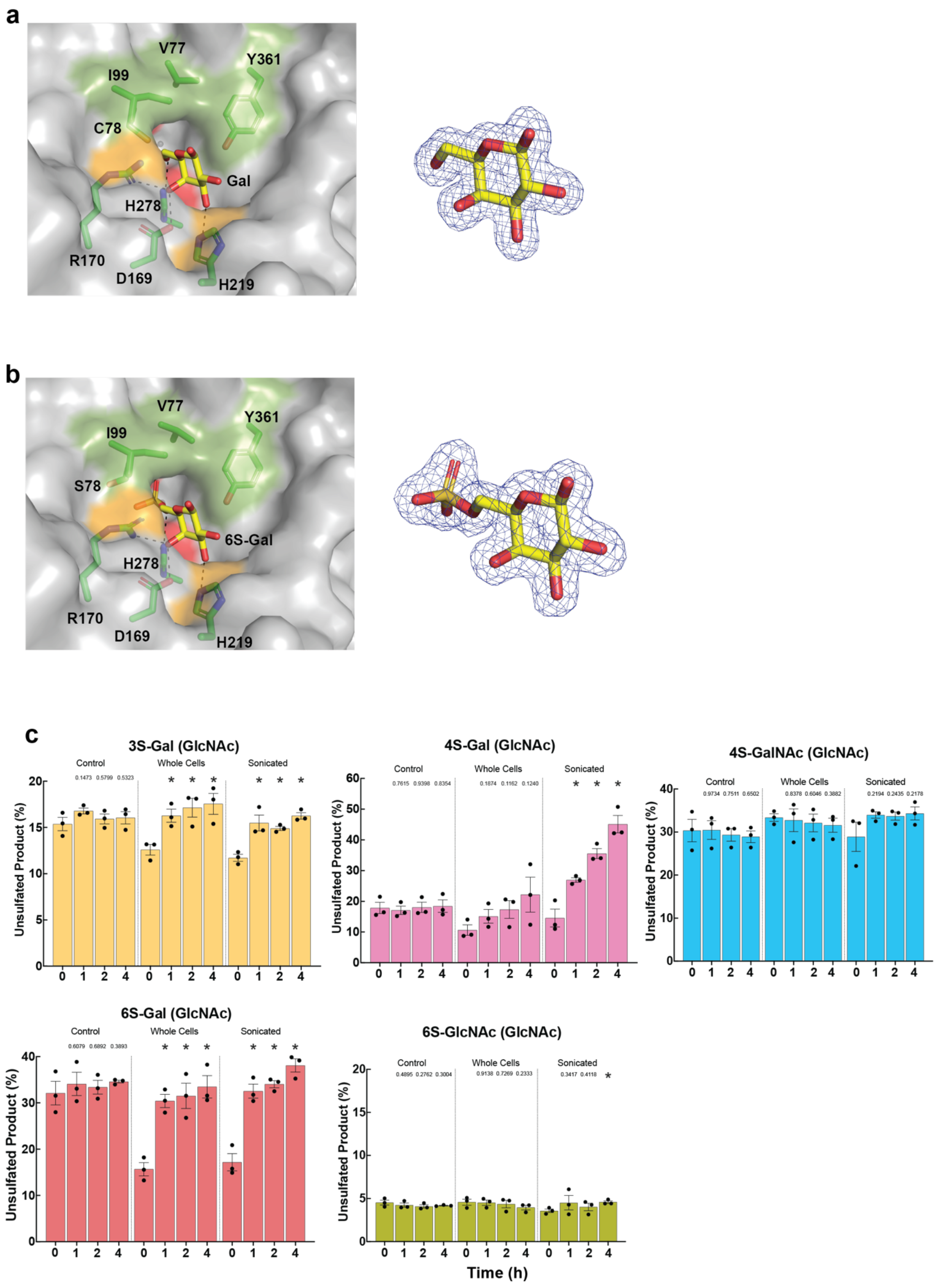


**Supplementary figure 4. Crystal structures and activity of Amuc0121$^{6S\text{-}Gal}$ and cellular localisation assays of *N*-acetyl-D-glucosamine grown *A. muciniphila* cells.**

**a, b.** Surface representation of the Amuc0121$^{6S\text{-}Gal}$ crystal structure in complex with D-galactose and *O*6 sulfated D-galactose, respectively. Key residues are shown as sticks beneath. In orange is the galactose recognition triad whilst green highlights the hydrophobic patch that accommodate Fuc (L-fucose); 6S-Gal is O6 sulfated D-galactose whilst GlcNAc is *N*-acetyl-D-glucosamine. In each case the the weighted 2mFobs-DFc electron density map, contoured at 1 $\sigma$, is shown on the right. **c.** HPAEC based cell assays with wildtype and transposon mutant sulfatases. Cells were grown with, *N*-acetyl-D-glucosamine as their carbon source, to mid-exponential phase before being left in aerobic conditions for ~30 mins to arrest growth. These were either left as whole cells or sonicated and 1 $\mu$M of a fluorescent substrates added to the media and sampled periodically. Experiments are biological triplicates. Statistical significance of time points versus control was calculated using an unpaired students t-test; an asterix indicates a value <0.5 otherwise the value is stated.

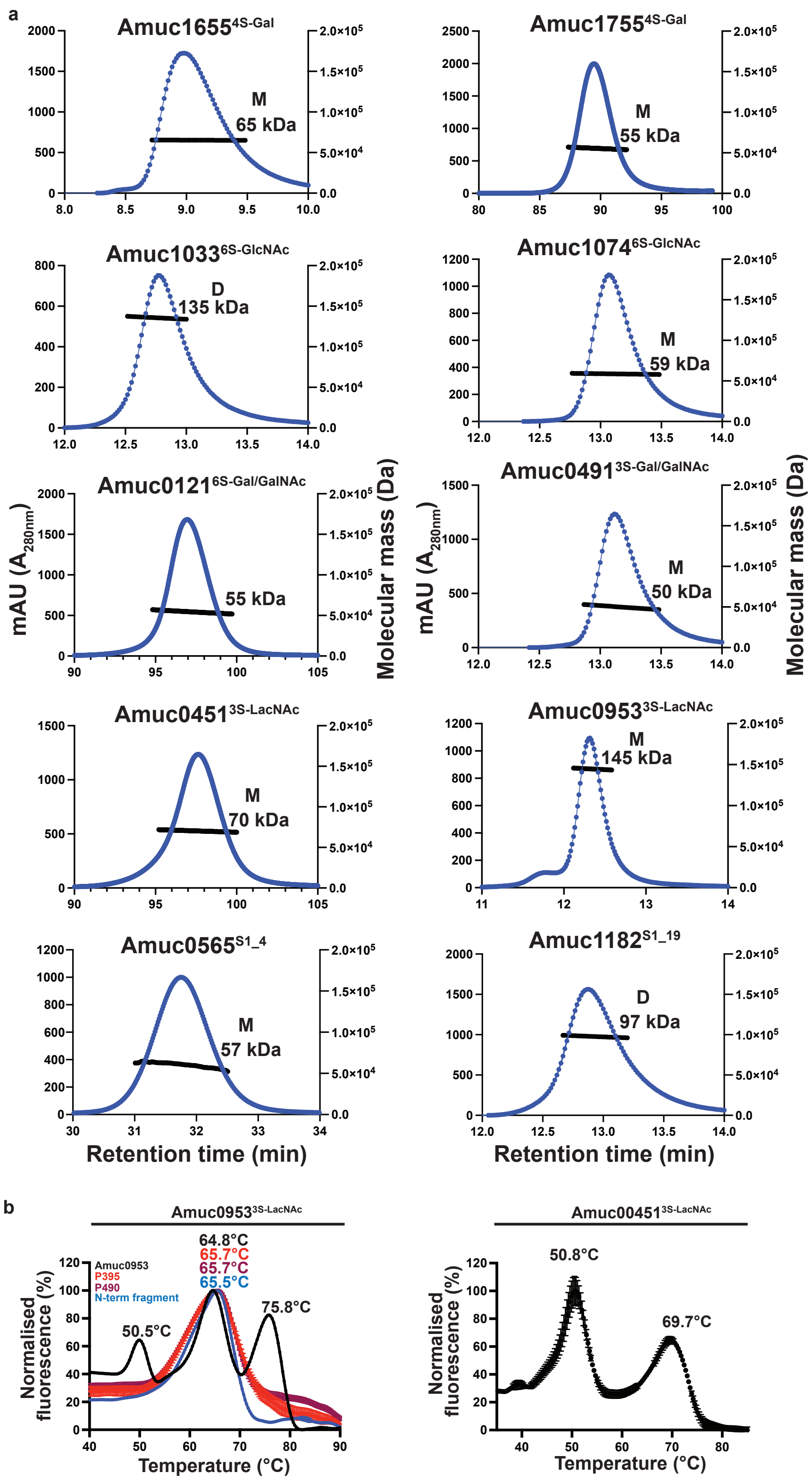


**Supplementary figure 5. Size determination and biophysical properties of *A. muciniphila* S1 sulfatases**

a. Size-exclusion chromatography coupled to light scattering profiles for *A. muciniphila* S1 sulfatases. In each case 5 mg/ml protein was utilised. **b.** Differential scanning fluorimetry profiles for Amuc0953$^{3S\text{-}LacNAc}$ and Amuc0953$^{3S\text{-}LacNAc}$ showing multiple, distinct unfolding events.

a

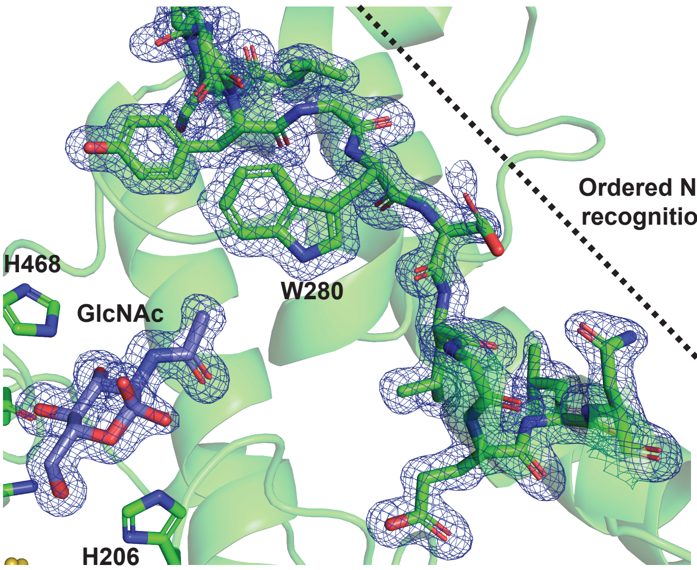


b

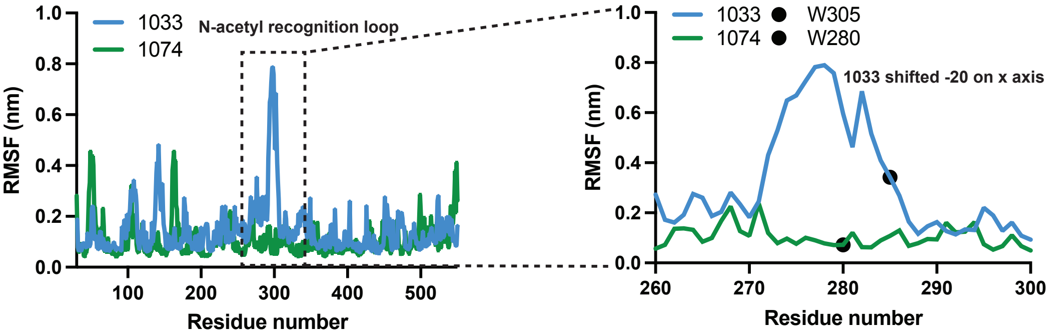


**Supplementary figure 6. Structure and dynamics of the S1_11 enzymes *N*-acetyl recognition loop structure.**

**a.** Cartoon and stick representation of the N-acetyl recognition loop of Amuc1074$^{6S\text{-}GlcNAc}$ in complex with GlcNAc; the weighted 2mFobs-DFc map has been contoured at 1 $\sigma$. **b.** Molecular dynamic simulations of unliganded Amuc1033$^{6S\text{-}GlcNAc}$ and Amuc1074$^{6S\text{-}GlcNAc}$ showing the R.M.S.F. values per residue with a zoomed in graph of the N-acetyl recognition loop.

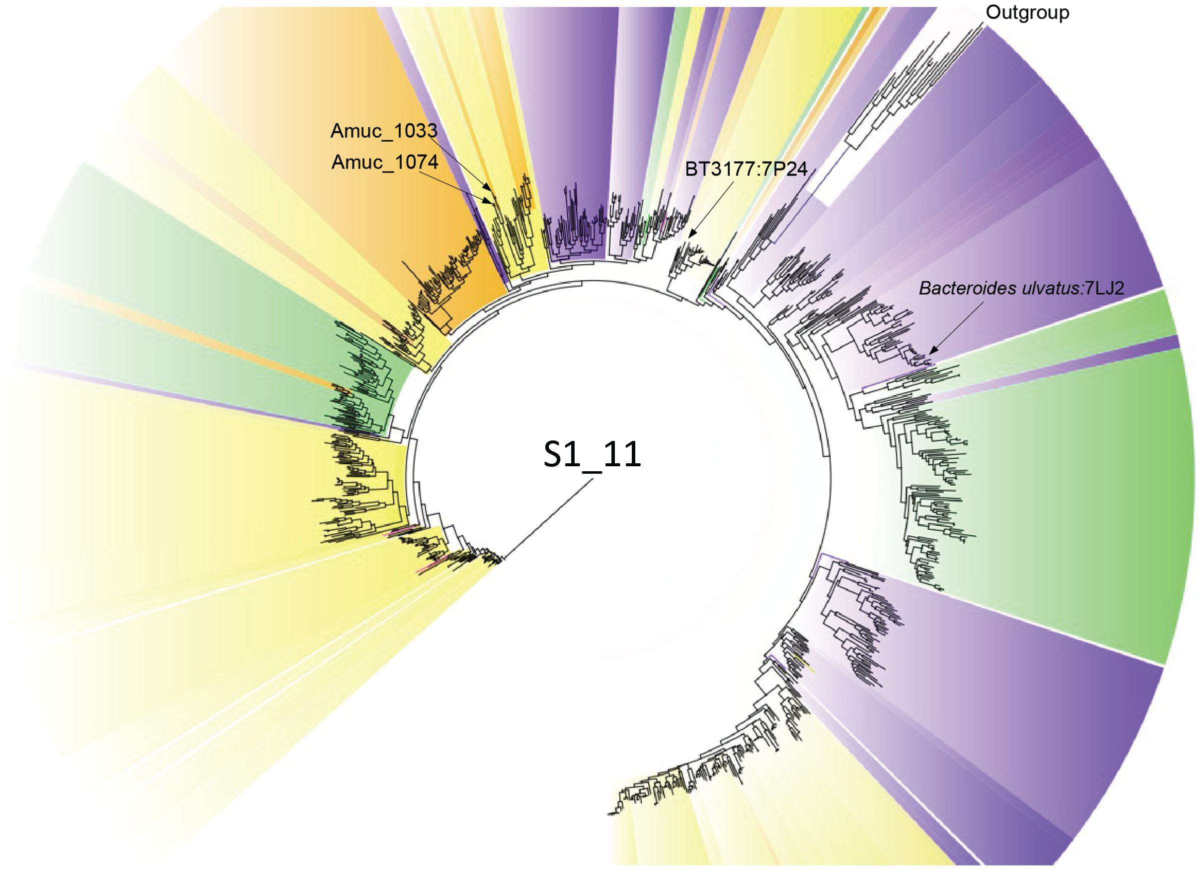


**Supplementary figure 7. Phylogenetic tree of the S1_11 subfamily**
Phylogenetic tree of 970 selected, most representative, sequences belonging to S1_11. The colours distinguish the sequences according to different lengths of the *N*-acetyl recognition loop, which in addition is highly variable in amino acid composition. Absence of loops and those with less than 5 amino acids are coloured in purple, those having 5 to 11 amino acids are coloured in green. Sequences having loops with lengths ranging from 12 to 19 and hydrophobic patches similar to Amuc1074 are coloured in yellow and those having loops longer than 19 amino acids (in general 20 to 25) are coloured in orange. Since the hydrophobic character (Trp in Amuc1074 and Amuc1033) is key for its interaction with the *N*-acetyl group of GlcNAc, some isolated sequences having loops without hydrophobic character are highlighted by pink lines.

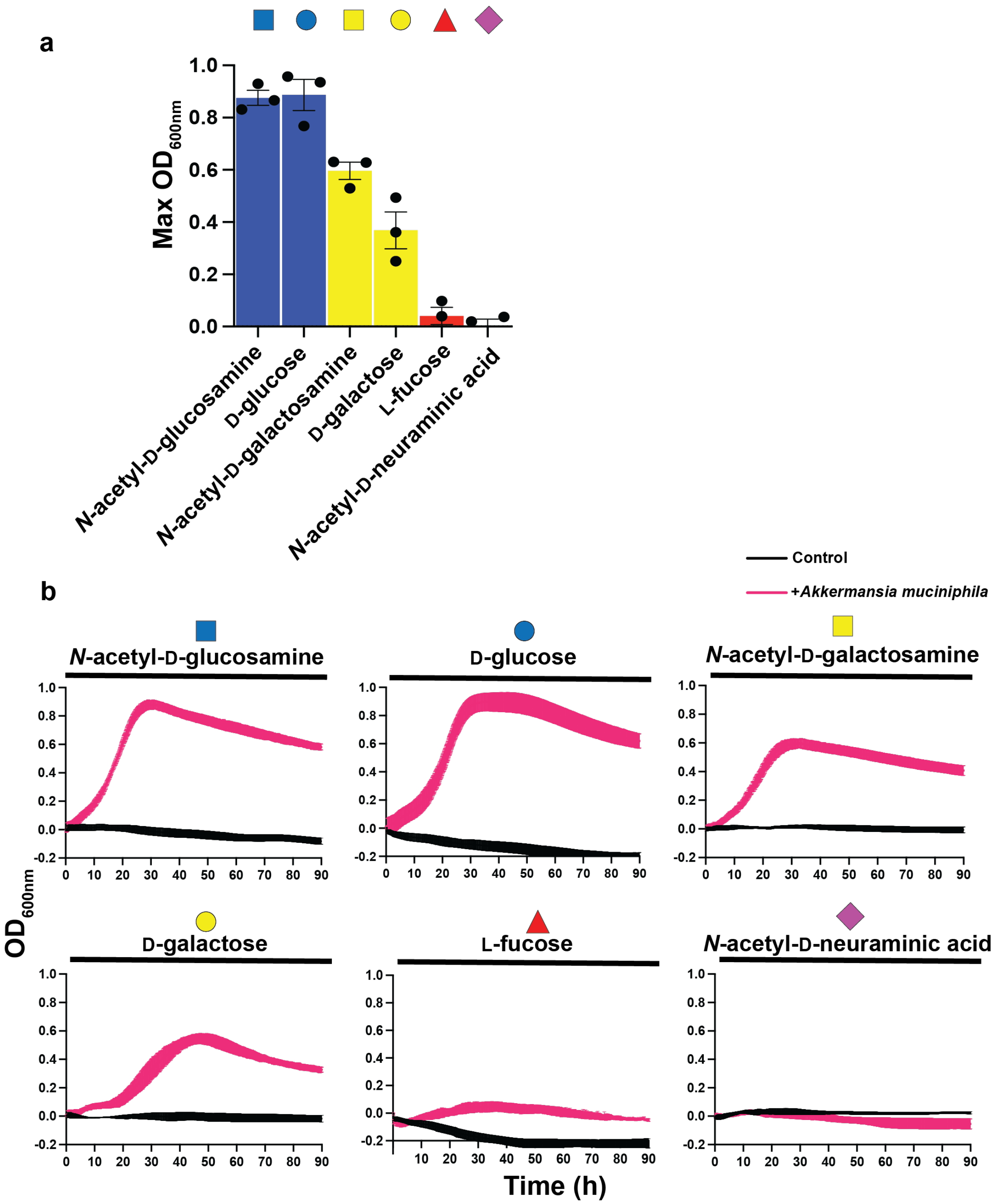


**Supplementary figure 8. Growth of *A. muciniphila* on monosaccharide growth substrates.**
**a.** Bar chart showing the maximum $OD_{600nm}$ each of the tested monosaccharides can allow *A. muciniphila* to reach **b.** Individual growth curves of *A. muciniphila* grown on monosaccharide substrates. All monosaccharides are at 10 mg/ml and the data are technical triplicates.

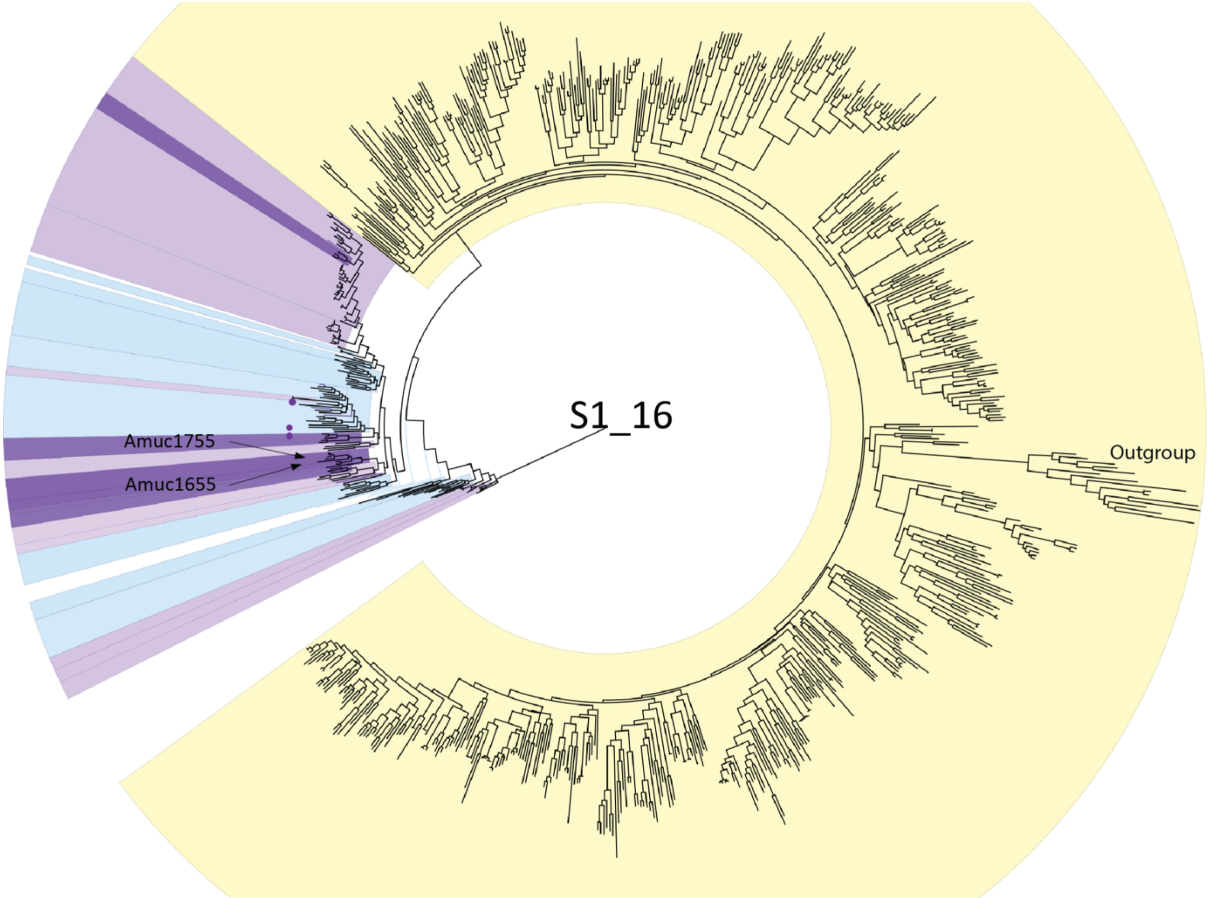


**Supplementary figure 9. Phylogenetic tree of the S1_16 subfamily**

Phylogenetic tree of 815 selected, most representative, sequences belonging to S1_16. Human and animal microbiome associated bacteria are coloured in light purple (mainly gut, faeces, and oral cavity), the dark purple are exclusively gut bacteria and are those sequences carrying the W_GEX stretch containing E464; light blue are aqueous environmental bacteria (i.e. waste water, rivulet, mangrove, Chinese sea sediment, etc). Notably, three sequences among these aquatic environments posses the W_GEX pattern (purple dots). All other sequences, coloured in yellow are bacteria isolated from various environmental, mainly soil samples.

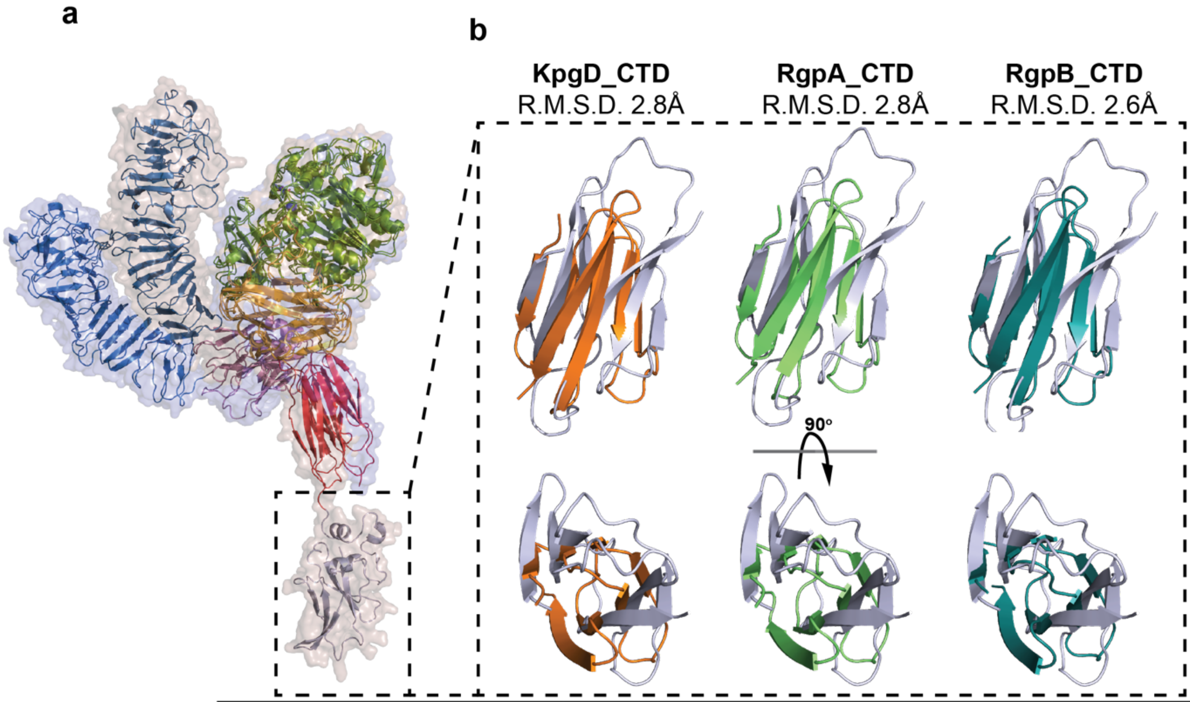


| | R.M.S.D (Å) | TM-score | Identity (%) | Aligned residues | Sequence length | Modelled residues |
|---|---|---|---|---|---|---|
| Amuc0953_CTD | - | - | - | - | 77 | 77 |
| Kgp_CTD | 2.8 | 0.41 | 12 | 50 | 63 | 63 |
| RgpA_CTD | 2.79 | 0.41 | 12 | 50 | 63 | 63 |
| RgpB_CTD | 2.64 | 0.43 | 12 | 50 | 63 | 63 |

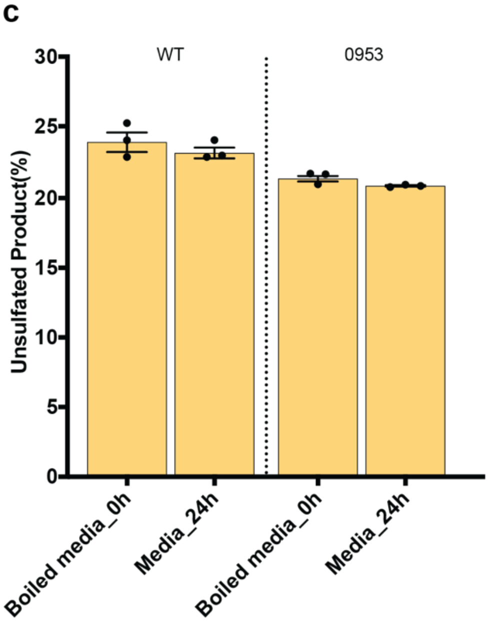


**Supplementary figure 10. Comparison of the Amuc0953$^{3S\text{-}LacNAc}$ crystal structure with its AF3 model, and 3S-Gal activity in the media.**

**a.** Overlay of the crystal structure of Amuc0953$^{3S\text{-}LacNAc}$ and its AF3 predicted model **b.** Comparison of the C-terminal domain (CTD) of Amuc0953$^{3S\text{-}LacNAc}$ with the CTD from type IX secretion systems. **c.** Filtered media from *A. muciniphila* cells grown on soluble porcine gastric mucin to mid exponential was tested for its ability to desulfate 3S-Gal.

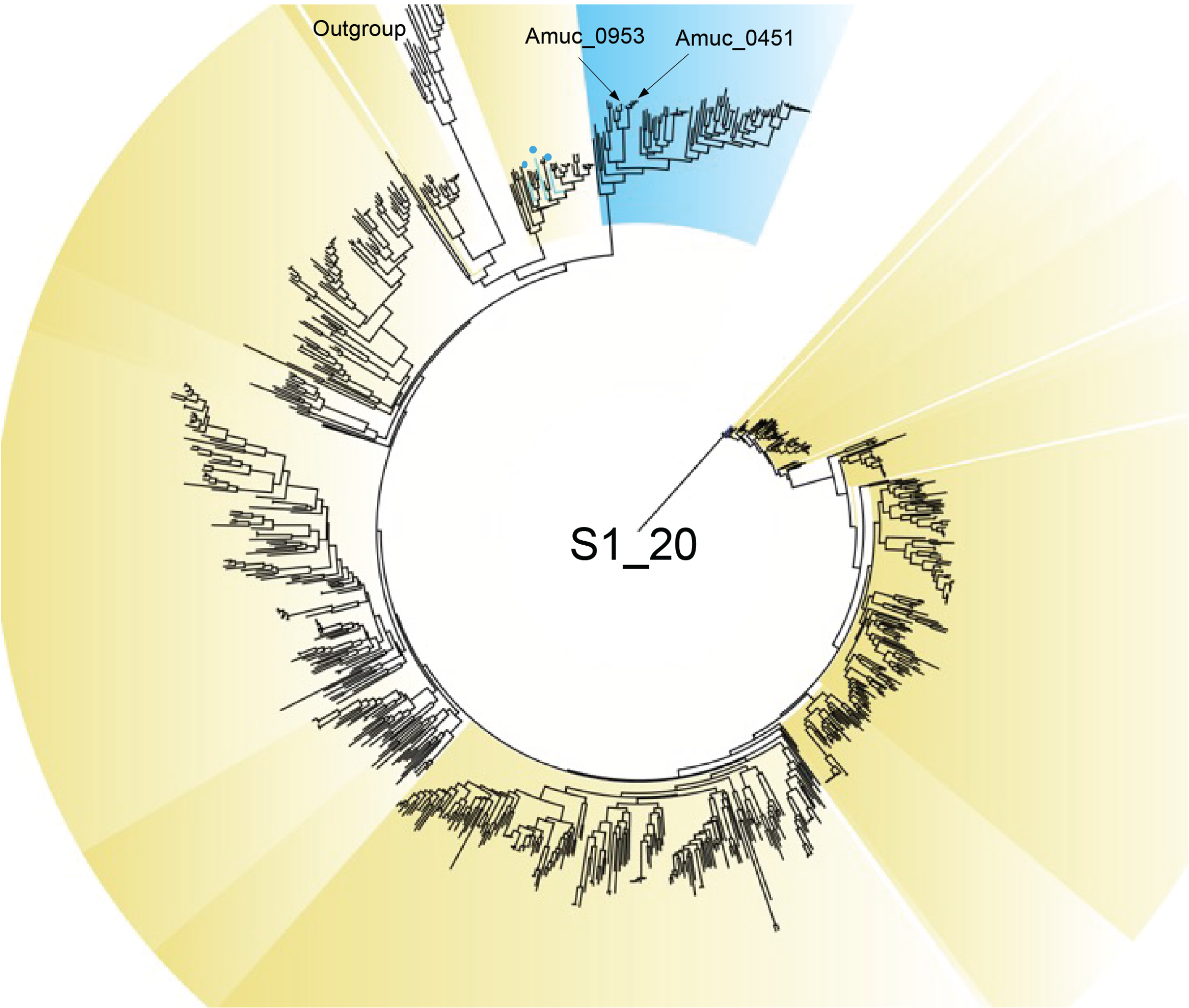


**Supplementary figure 11. Phylogenetic tree of the S1_20 subfamily**

Phylogenetic tree of 848 selected, most representative, sequences belonging to S1_20. The sequences carrying the concomitant Gln/His & Glu/Phe mutations are highlighted. Three sequences from Streptomyces sp. among the neighbouring group, all belonging to the Actinomycetota phylum, also contain the Gln/His & Glu/Phe mutations and are marked by blue lines and dots. The sequences of the branch highlighted in blue all belong either to the Bacteroidota, Planctomycetota or Verrucomicrobiota phylum.

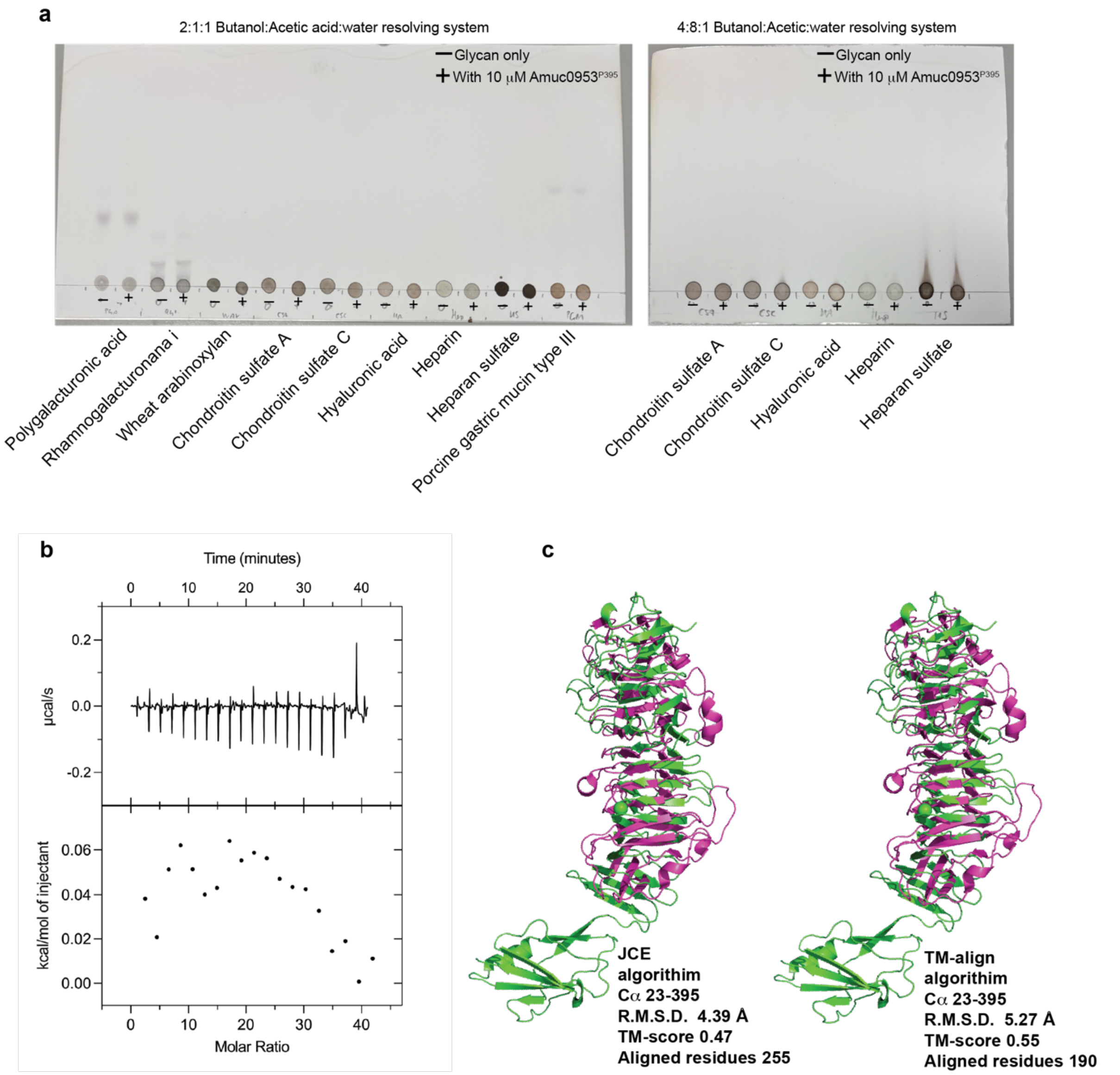


**Supplementary figure 12. Testing of Amuc0953$^{P395}$ for catalytic and bidnding activity against selected host glycans, and comparison to CBM89.**

**a.** Thin layer chromatography of various polysaccharides incubated with Amuc0953$^{P395}$. For highly charged and sulfated glycans a second solvent system was used to ensure that any oligosaccharide products produced had the ability to migrate. All reactions were in 10 mM HEPES pH 7.0, with 150 mM NaCl, and 10 μM Amuc0953$^{P395}$. **b.** Isothermal titration calorimetry of 50 μM Amuc0953$^{P395}$ against 5 mg/ml wheat arabinoxylan. **c.** structural overlay of Amuc0953$^{P395}$ with CBM89 (PBMDCECB_09513; PDB:7JIV), the only other CBM to have a β-helix fold. **d.** Comparison of the area housing the key binding residues in Amuc0953$^{P395}$ and CBM89 (PBMDCECB_09513) from the JCE algorithm overlay.

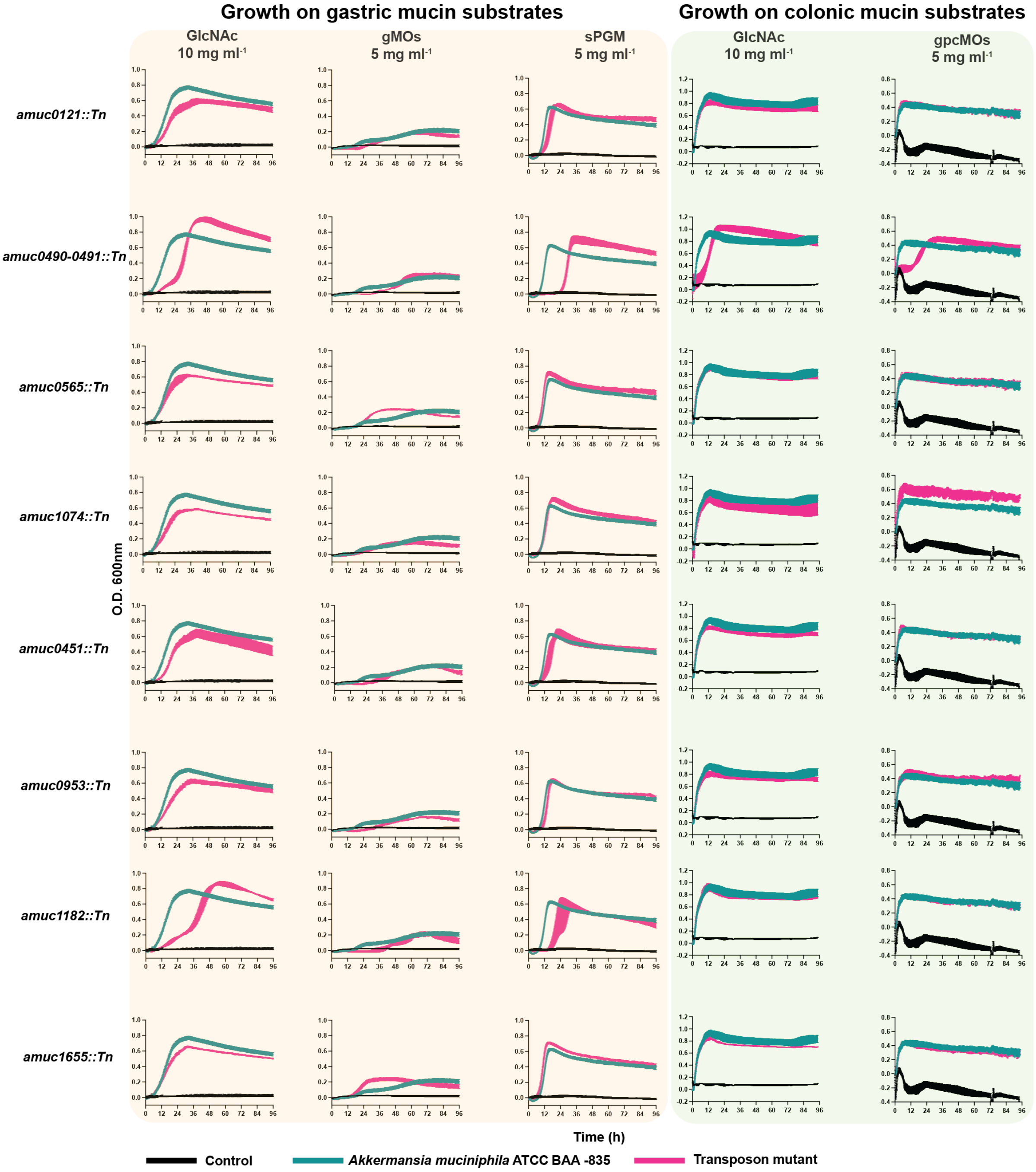


**Supplementary figure 13. Growth of transposon mutants on mucin substrates.** Bacterial growth curves of *Akkermansia muciniphila* ATCC BAA-835 and sulfatase transposon mutants on gastric and colonic mucin derived substrates. All growth curves are technical triplicates performed in CMM as a base media supplemented with 5 mg ml$^{-1}$ of the appropriate carbon source as listed. GlcNAc = *N*-acetyl D-glucosamine, gMOs = gastric mucin oligosaccharides, sPGM = soluble porcine gastric mucin, and gpcMOs = glycoprotein porcine colonic mucin oligosaccharides.

**Supplemental table 1. Kinetic rates of *A. muciniphila* against fluorescently linked carbohydrates**

All experiments were performed in technical triplicate in a buffer (100 mM) at the appropriate pH optimum supplemented with 150 mM NaCl and $CaCl_2$. NQ indicates the enzyme is active but the activity could not be accurately quantified; NA indicates the enzyme is not active on the substrate; NT indicates the enzyme was not tested quantitatively against the substrate. Values are $k_{cat}/K_M$ in units of $min^{-1}$ $M^{-1}$.

| | 4S-Gal | 4S-GalNAc | 6S-GlcNAc | 6S-GlcNS | 3S-Gal | 3S-GalNAc | 3S-LacNAc | 3S-Lac-N-biose | 3S-lewisX | 3S-lewisA | 6S-Gal | 6S-GalNAc | +10mM EDTA |
|---|---|---|---|---|---|---|---|---|---|---|---|---|---|
| Amuc1755 | (9.73 ± 0.28) x10$^{2}$ | NQ | NA | NA | NA | NA | NA | NA | NA | NA | NA | NA | (2.71 ± 0.11) x10$^{2}$ |
| Amuc1755$^{E464G}$ | NQ | (2.35 ± 0.65) x10$^{2}$ | NA | NA | NA | NA | NA | NA | NA | NA | NA | NA | NT |
| Amuc1755$^{E464A}$ | NQ | (2.00 ± 0.61) x10$^{2}$ | NA | NA | NA | NA | NA | NA | NA | NA | NA | NA | NT |
| Amuc1655 | (7.43 ± 0.18) x10$^{1}$ | NQ | NA | NA | NA | NA | NA | NA | NA | NA | NA | NA | (6.34 ± 0.06) x10$^{1}$ |
| Amuc1074 | NA | NA | (2.01 ± 0.03) x10$^{3}$ | (2.43 ± 0.02) x10$^{3}$ | NA | NA | NA | NA | NA | NA | NA | NA | (1.90 ± 0.03) x10$^{3}$ |
| Amuc1033 | NA | NA | (1.28 ± 0.02) x10$^{3}$ | (1.41 ± 0.06) x10$^{3}$ | NA | NA | NA | NA | NA | NA | NA | NA | (1.07 ± 0.06) x10$^{3}$ |
| Amuc0121 | NA | NA | NA | NA | NA | NA | NA | NA | NA | NA | NQ | NQ | |
| Amuc0491 | NA | NA | NA | NA | (1.04 ± 0.03) x10$^{2}$ | (1.77 ± 0.05) x10$^{3}$ | (2.21 ± 0.14) x10$^{2}$ | (9.17 ± 0.04) x10$^{2}$ | (2.66 ± 0.06) x10$^{1}$ | (2.93 ± 0.06) x10$^{1}$ | NA | NA | (3.69 ± 0.20) x10$^{2}$ |
| Amuc0451 | NA | NA | NA | NA | (3.89 ± 0.19) x10$^{2}$ | NA | (3.50 ± 0.11) x10$^{3}$ | (2.50 ± 0.06) x10$^{3}$ | (2.92 ± 0.06) x10$^{1}$ | (1.09 ± 0.02) x10$^{2}$ | NA | NA | (2.41 ± 0.03) x10$^{3}$ |
| Amuc0953 | NA | NA | NA | NA | (2.14 ± 0.2) x10$^{2}$ | NA | (4.83 ± 0.15) x10$^{3}$ | (3.84 ± 0.07) x10$^{3}$ | NT | NT | NA | NA | (4.10 ± 0.02) x10$^{3}$ |
| BT1636 | NA | NA | NA | NA | (8.12 ± 0.13) x10$^{3}$ | NA | (1.61 ± 0.13) x10$^{4}$ | NT | NT | NT | NA | NA | NT |

**Supplemental table 2. Cell viability counting**

Total cell counts, and the percentage of viable and nonviable (dead or dying) cells observed for exponential phase *A. muciniphila* cells incubated in chopped meat broth (CMB) or phosphate buffered saline (PBS). The number of viable cells in the exponential phase *starter* culture and the sample incubated for 4 hours in CMB represented >99 % of the total cell population. Incubation of the *A. muciniphila* cells in PBS for 4 hours greatly increased the number of cells with compromised membranes. In PBS, we observed two populations of propidium iodine-stained cells with different levels of fluorescence emission (intense or dim). All cells in these two populations were also visible in the SYTO-9 fluorescence channel. The two populations were assigned to cells with high (intense florescence, 16.6%) and low (dim fluorescence, 17.4%) levels of membrane damage, and they were combined for the purpose of calculating the number of nonviable cells in PBS. The data presented for each condition represent combined cell counts from three separate biological replicates (n = 3).

| **Condition** | **Total number of cells counted** | **Viable cells (%)** | **Nonviable cells (%)** | **Notes** |
|---|---|---|---|---|
| *A. muciniphila* exponential phase *starter* culture grown in CMM | 3560 | 99.7 | 0.337 | Staining with SYTO-9 and propidium iodide was done in CMB. |
| *A. muciniphila* cells washed and incubated in CMM for 4 hours | 5217 | 99.8 | 0.249 | Staining with SYTO-9 and propidium iodide was done in CMB. |
| *A. muciniphila* cells washed and incubated in PBS for 4 hours | 1105 | 66.1 | 33.9 | Staining with SYTO-9 and propidium iodide was done in PBS.<br>**16.6%** and **17.4%** of the nonviable cells were intensely and dimly fluorescent, respectively, after staining with propidium iodide. |
| CMB = chopped meat broth, PBS = phosphate buffered saline pH 7.4<br>Cells were stained with the following final dye concentrations in the viability assay: 5 μM SYTO-9 and 2 μM propidium iodide. | | | | |

**Supplemental table 3. Primers used for cloning**

| | Cloning Primers | Primer sequence (3' to 5')& |
|---|---|---|
| **Amuc_0121** | Amuc_0121_Fw_Ntag | CATCATCACCACCATCACGAGAACCTGTACTTCCAGGGCacgccaacggtcaagc |
| **Amuc_0121** | Amuc_0121_Rv_Ntag | GTGGCGGCCGCTCTATTAgtcgtccaagggaagcag |
| **Amuc_0451** | Amuc_0451_Fw_Ntag | CATCATCACCACCATCACGAGAACCTGTACTTCCAGGGCgcgaagccggctgcc |
| **Amuc_0451** | Amuc_0451_Rv_Ntag | GTGGCGGCCGCTCTATTAcctggattccttcccgtg |
| **Amuc_0491** | Amuc_0491_Fw_Ntag | CATCATCACCACCATCACGAGAACCTGTACTTCCAGGGCgcacggcccaatgtg |
| **Amuc_0491** | Amuc_0491_Rv_Ntag | GTGGCGGCCGCTCTATTAttcggctacctgttccg |
| **Amuc_0565** | Amuc_0565_Fw_Ntag | CATCATCACCACCATCACGAGAACCTGTACTTCCAGGGCgacaggcctaatatcgtc |
| **Amuc_0565** | Amuc_0565_Rv_Ntag | GTGGCGGCCGCTCTATTAgggcgcgtatgcccgtttttctgc |
| **Amuc_0953** | Amuc_0953_Fw_Ntag | CATCATCACCACCATCACGAGAACCTGTACTTCCAGGGCgctgatgttgaatatgtc |
| **Amuc_0953** | Amuc_0953_Rv_Ntag | GTGGCGGCCGCTCTATTAtctcgtcacctttaatctcagg |
| **Amuc_0953** | Amuc_0953_Ndom_Stop_Fw | gacgctgcagacggatgta**TAA**gcggaaaagccggacatgc |
| **Amuc_0953** | Amuc_0953_Ndom_Stop_Rv | gcatgtccggcttttccgc**TTA**tacatccgtctgcagcgtc |
| **Amuc_0953** | Amuc_0953_P395Stop_Fw | ggggatatttccctgccc**TAA**agttcatccctgacgctgg |
| **Amuc_0953** | Amuc_0953_P395Stop_Rv | ccagcgtcagggatgaact**TTA**gggcagggaaatatcccC |
| **Amuc_1033** | Amuc_1033_Fw_Ntag | CATCATCACCACCATCACGAGAACCTGTACTTCCAGGGCgaccagcctcaaaag |
| **Amuc_1033** | Amuc_1033_Rv_Ntag | GTGGCGGCCGCTCTATTAatctctggagggagcg |
| **Amuc_1074** | Amuc_1074_Fw_Ntag | CATCATCACCACCATCACGAGAACCTGTACTTCCAGGGCcaaaccaaggctgag |
| **Amuc_1074** | Amuc_1074_Rv_Ntag | GTGGCGGCCGCTCTATTAccaggaagcgtcgaatttggg |
| **Amuc_1182** | Amuc_1182_Fw_Ntag | CATCATCACCACCATCACGAGAACCTGTACTTCCAGGGCcagccccgcctttcc |
| **Amuc_1182** | Amuc_1182_Rv_Ntag | GTGGCGGCCGCTCTATTActcaggctgcctgtccgg |
| **Amuc_1655** | Amuc_1655_Fw_Ntag | CATCATCACCACCATCACGAGAACCTGTACTTCCAGGGCaatggttccaacactgtttc |
| **Amuc_1655** | Amuc_1655_Rv_Ntag | GTGGCGGCCGCTCTATTAaaaaccattccccttcacctgg |
| **Amuc_1755** | Amuc_1755_Fw_Ntag | CATCATCACCACCATCACGAGAACCTGTACTTCCAGGGCgcttccgtaaaagcat |
| **Amuc_1755** | Amuc_1755_Rv_Ntag | GTGGCGGCCGCTCTATTAcagcctggccgccg |

**Supplemental table 4. Primers used for mutagenesis**

| | Mutagenesis Primers | Primer sequence (5' to 3') |
|---|---|---|
| **Amuc_0121** | Cys78Ser_Fw | ACATCCGTCTCTACCCCTTCCCGCTATGCCCTGTTC |
| **Amuc_0121** | Cys78Ser_Rv | GGAAGGGGAGAAGACGGATGTGGTGGAATAGGCGTC |
| **Amuc_1755** | Glu464Ala_Fw | GCCGGGAACGGAAACAACTCCCTTTAT |
| **Amuc_1755** | Glu464Ala_Rv | CCCCCATACGTTGGGCGTGTGGAA |
| **Amuc_1755** | Glu464Gly_Fw | GGCGGGAACGGAAACAACTCCCTTTAT |
| **Amuc_1755** | Glu464Gly_Rv | CCCCCATACGTTGGGCGTGTGGAA |

### Supplemental table 5. Chopped meat growth media growth composition

| **Components** | **Amount (per 250 ml)** |
|---|---|
| Beef Extract | 2.5g |
| Pancreatic Digest of Casein | 7.5g |
| Yeast Extract | 1.25g |
| Potassium Phosphate Dibasic ($K_2HPO_4$) | 1.25g |
| Cysteine | 250mg |
| Vitamin K (1mg/ml) | 250µl |
| Vitamin B3 (0.01mg/ml) | 250µl |
| Hematin (1.2mg/ml)-Histidine (0.2M) solution | 1ml |
| Balch's Vitamins (0.04mg/ml) | 2.5ml |
| Trace Mineral Solution (5.35mg/ml) | 2.5ml |
| Purine/Pyrimidine Solution (1mg/ml) | 2.5ml |
| Amino Acid Solution (5mg/ml) | 2.5ml |
| **Adjust pH to 7.2 and filter sterilise** | |
| **Amino Acid solution**<br>Alanine, Arginine, Asparagine, Aspartic Acid, Cysteine, Glutamic Acid, Glutamine, Glycine , Histidine, Isoleucine, Leucine, Lysine, Methionine, Phenylalanine, Proline, Serine, Threonine, Tryptophan, Tyrosine, Valine<br><br>Dissolve 62.5mg each of the 20 essential amino acid to 250 ml (5mg/ml total stock). Filter sterilize, store at room temperature. | |
| **Balch's Vitamins**<br>*p*-Aminobenzoic acid, 5mg<br>Folic acid, 2mg (Sigma, F7876)<br>Biotin, 2mg (Sigma, B4501)<br>Nicotinic acid, 5mg (Sigma, N4126)<br>Calcium pantothenate, 5mg (Sigma, P2250)<br>Riboflavin, 5mg (Sigma, R7649)<br>Thiamine HCl, 5mg (Sigma, T4625)<br>Pyridoxine HCl (vitamin $B_6$), 10mg<br>Cyanocobalamin (vitamin $B_{12}$), 0.1mg<br>Thioctic acid (lipoic acid), 5mg<br>Distilled Water, 1L<br><br>Dissolve vitamins, pH to 7.0, filter sterilize, keep at 4C in dark. | |
| **Trace mineral supplement**<br>EDTA, 0.5g (Sigma, ED4SS)<br>$MgSO_4*7H_2O$, 3g<br>$MnSO_4*H_2O$, 0.5g<br>NaCl, 1g (Sigma, S7653)<br>$FeSO_4*7H_2O$, 0.1g (Sigma, 215422)<br>$CaCl_2$, 0.1g<br>$ZnSO_4*7H_2O$, 0.1g<br>$CuSO_4*5H_2O$, 0.01g<br>$H_3BO_3$, 0.01g (Sigma, B6768)<br>$Na_2MoO_4*2H_2O$, 0.01g<br>$NiCl_2*6H_2O$, 0.02g<br><br>Dissolve in 1L, pH to 7.0, filter sterilize and store at room temperature. | |
| **Purine/Pyrimidine Solution**<br>Adenine (Sigma, A2786) ,Guanine (Sigma, G11950), Thymine (Sigma, T0895)<br>Cytosine (Sigma, C3506), Uracil (Sigma, U1128)<br><br>Dissolve 200mg each into 1L (1mg/ml total stock), pH to7.0, filter sterilize and store at room temperature | |

**Supplemental table 6. Minimal media for *Bacteroides thetaiotaomicron* growths**

| stock | 100 ml | |
|---|---|---|
| $NH_4$ $SO_4$ | 0.1 | g |
| $Na_2CO_3$ | 0.1 | g |
| cysteine, free base | 0.05 | g |
| 1 M $KPO_4$ pH 7.2 | 10 | ml |
| Vitamin K solution, 1mg/ml | 0.1 | ml |
| $FeSO_4$, 0.4 mg/ml | 1 | ml |
| resazurin, 0.25 mg/ml | 0.4 | ml |
| Vitamin $B_{12}$, 0.01 mg/ml | 0.05 | ml |
| Mineral Salts for defined medium | 5 | ml |
| 1.2 mg/ml Haematin in 0.2 M histidine pH 8.0 | 0.1 | ml |
| $dH_2O$ | 85 | ml |

Mineral salts for defined media:

NaCl 18 g
$CaCl_2$ $2H_2O$ 0.53 g
$MgCl_2$ $6H_2O$ 0.40 g
$MnCl_2$ $4H_2O$ 0.20 g
$CoCl_2$ $6H_2O$ 0.20g

Dissolve in 1 litre and filter; 0.2 micron.

**Supplemental table 7. Crystallography collection and refinement statistics**

Values in parenthesis are for the highest resolution shell. $R_{free}$ was calculated using a set (5%) of randomly selected reflections that were excluded from refinement.

| **Collection statistics** | Amuc0121$^{6S\text{-}Gal}$ Gal | Amuc0121$^{6S\text{-}Gal}$ 6S-Gal | Amuc0121$^{6S\text{-}Gal}$ 6'S-LewisA | Amuc0451$^{3S\text{-}Gal}$ Apo |
|---|---|---|---|---|
| Wavelength (Å) | 1.00 | 0.98 | 0.98 | 1.00 |
| Resolution (Å) | 61.53-1.56 (1.59-1.56) | 60.27—1.83 (1.83-1.80) | 58.75-1.92 (1.95-1.92) | 59.90-1.60 (1.63-1.60) |
| Space group | $P2_12_12_1$ | $P2_12_12_1$ | $P2_1$ | $P2_1$ |
| Unit-cell parameters | | | | |
| a, b, c (Å) | 66.510, 91.490 162.080 | 65.056,91.127 160.101 | 135.040,64.150 151.360 | 113.660 59.900 123.720 |
| α, β, γ (°) | 90, 90, 90 | 90, 90, 90 | 90.0,104.88 90.0 | 90.000 115.75 90.000 |
| No. of measured reflections | 914204 (37811) | 1210143 (61485) | 1313104 (56063) | 1291159 (55288) |
| No. of independent reflections | 141184 (6959) | 89016 (4487) | 191925 (8643) | 197802 (9822) |
| Completeness (%) | 100 (99.9) | 100 (100) | 99.4 (90.7) | 100 (100) |
| Redundancy | 6.5 (5.4) | 13.6 (13.7) | 6.8 (6.5) | 6.5 (5.6) |
| <I>/<σ(I)> | 15.8 (1.4) | 8.0 (0.8) | 9.3 (0.8) | 10.3 (1.4) |
| CC(1/2) | 0.999 (0.589) | 0.996 (0.446) | 0.991 (0.421) | 0.998 (0.549 |
| **Refinement statistics*** | | | | |
| $R_{work}/R_{free}$ (%) | 14/18 | 18/22 | 21/24 | 15/19 |
| No. of non-H atoms | 15878 | 15449 | 38424 | 22414 |
| No. of protein, atoms | 15216 | 15089 | 37797 | 21234 |
| No. of solvent atoms | 596 | 312 | 363 | 951 |
| No. of ligand atoms | 40 | 48 | 264 | 230 |
| r.m.s. deviation from ideal values values | | | | |
| Bond angle (°) | 1.63 | 1.58 | 1.56 | 1.67 |
| Bond length (Å) | 0.01 | 0.007 | 0.007 | 0.008 |
| Average B factor (Å$^2$) | | | | |
| Protein (by chain) | 27.7/34.5 | 25.2/26.7 | 34.8/51.9/42.2/ 38.5/33.8 | 23.2/23.1 |
| Solvent | 37.7 | 24.3 | 31.2 | 35.6 |
| Ligand (by chain) | 18.9/21.7 | 16.9/18.3 | 29.4/62.9/40.4/ 54.0/33.6 | N/A |
| Ramachandran plot$^+$, most favoured regions (%) | 96.2 | 96.2 | 96.4 | 97.0 |
| Molprobity score | 1.28 | 1.46 | 1.57 | 1.22 |
| PDB code | **9S7X pdb_00009S7X** | **9S85 pdb_00009S85** | **9S88 pdb_00009S88** | **9S8A pdb_00009S8A** |

**Supplemental table 8. Crystallography collection and refinement statistics**
Values in parenthesis are for the highest resolution shell. $R_{free}$ was calculated using a set (5%) of randomly selected reflections that were excluded from refinement.

| **Collection statistics** | Amuc0953$^{P490}$ Apo | Amuc0953$^{3S\text{-}LacNAc}$ Gal | Amuc1074$^{6S\text{-}GlcNAcl}$ GlcNAc | Amuc1755$^{4S\text{-}Gal}$ Gal |
|---|---|---|---|---|
| Wavelength (Å) | 0.8 | 1.0 | 0.98 | 1.0 |
| Resolution (Å) | 55.34-2.00 (2.04-2.00) | 78.35-2.90 (2.95-2.90) | 71.88-1.38 (1.40-1.38) | 71.54-2.10 (2.16-2.10) |
| Space group | $P2_1$ | $I4_1$ | $I4_1$ | $P6_4$ |
| Unit-cell parameters | | | | |
| a, b, c (Å) | 100.44, 59.15, 111.12 | 187.280 187.280 221.770 | 130.01 ,130.01 86.27 | 143.08 143.08 50.41 |
| α, β, γ (°) | 90, 95.11, 90 | 90,90,90 | 90,90,90 | 90,90,120 |
| No. of measured reflections | 606234 (31455) | 1156298 (61502) | 990308 (50221) | 344730 (28425) |
| No. of independent reflections | 88190 (4512) | 84317 (4457) | 146947 (7245) | 34749 (2818) |
| Completeness (%) | 99.99 (99.99) | 100 (100) | 100 (100) | 100 (100) |
| Redundancy | 6.9 (7.0) | 13.7 (13.8) | 6.7 (6.9) | 9.9 (10.1) |
| <I>/<σ(I)> | 6.6 (1.2) | 9.9 (1.0) | 10.7 (1.4) | 7.7 (1.6) |
| CC(1/2) | 0.994 (0.364) | 0.998 (0.641) | 0.998 (0.707) | 0.994 (0.666) |
| **Refinement statistics*** | | | | |
| $R_{work}/R_{free}$ (%) | 19/23 | 21/26 | 17/18 | 17/21 |
| No. of non-H atoms | 13662 | 36806 | 9101 | 8465 |
| No. of protein, atoms | 13320 | 36762 | 8618 | 8330 |
| No. of solvent atoms | 265 | N/A | 437 | 116 |
| No. of ligand atoms | N/A | 19 | 46 | 19 |
| r.m.s. deviation from ideal values | | | | |
| Bond angle (°) | 0.007 | 0.006 | 0.011 | 0.007 |
| Bond length (Å) | 1.56 | 1.56 | 1.90 | 1.73 |
| Average B factor (Å$^2$) | | | | |
| Protein (by chain) | 49.3/41.7 | 66.5/66.4 | 23.3 | 37.7 |
| Solvent | 43 | N/A | 30.8 | 34.3 |
| Ligand (by chain) | N/A | 75.5/98.1 | 16.8 | 59.5 |
| Ramachandran plot$^+$, most favoured regions (%) | 96.8 | 90.4 | 96.2 | 96.0 |
| Molprobity score | 1.50 | 2.28 | 1.37 | 1.83 |
| PDB code | **9SF6 pdb_00009SF6** | **9S8X pdb_00009S8X** | **9S8Y pdb_00009S8Y** | **9S8F pdb_00009S8F** |
